# A low-level perceptual correlate of behavioral and clinical deficits in ADHD

**DOI:** 10.1101/199216

**Authors:** Andra Mihali, Allison G Young, Lenard A. Adler, Michael M. Halassa, Wei Ji Ma

**Author notes:** Corresponding authors are Wei Ji Ma and Michael M Halassa. Andra Mihali and Allison G Young contributed equally to this work.

## Abstract

In many studies of attention-deficit hyperactivity disorder (ADHD), stimulus encoding and processing (perceptual function) and response selection (executive function) have been intertwined. To dissociate deficits in these functions, we introduced a task that parametrically varied low-level stimulus features (orientation and color) for fine-grained analysis of perceptual function. It also required participants to switch their attention between feature dimensions on a trial-by-trial basis, thus taxing executive processes. Furthermore, we used a response paradigm that captured task-irrelevant motor output (TIMO), reflecting failures to use the correct stimulus-response rule. ADHD participants had substantially higher perceptual variability than Controls, especially for orientation, as well as higher TIMO. In both ADHD and Controls, TIMO was strongly affected by the switch manipulation. Across participants, the perceptual variability parameter was correlated with TIMO, suggesting that perceptual deficits are associated with executive function deficits. Based on perceptual variability alone, we were able to classify participants into ADHD and Controls with a mean accuracy of about 77%. Participants’ self-reported General Executive Composite score correlated not only with TIMO but also with the perceptual variability parameter. Our results highlight the role of perceptual deficits in ADHD and the usefulness of computational modeling of behavior in dissociating perceptual from executive processes.

## Introduction

In Attention Deficit Hyperactivity Disorder (ADHD), the behavioral deficits captured by self-reports and collateral reports have been attributed to differences in attention, executive function, and lower-level processes, including perceptual function. In the realm of visual attention, differences in accuracy or reaction time have been found in some visual search tasks but not in others (for a review, see [1]). No consistent deficits have been found when probing selective attention with visuo-spatial orienting tasks [2–5]. ADHD patients tend to have worse executive function than Controls [6–9], predominantly in response execution and inhibition [10–12], but also in working memory and switching between stimulus-response rules [13–16]. While some researchers believe executive function impairments to be primary in ADHD, others acknowledge that they are neither necessary nor sufficient to cause the disorder [6, 7]. More specifically, yet others suggest that ADHD impairments are a combination of deficits in high-level and “low-level processes” [17–22]. These low-level processes entail arousal [23], relatedly, accumulation of evidence [24], timing [25], or reward sensitivity (see [26, 27] for reviews). It should be kept in mind that ADHD might be a heterogenous disorder [28, 29] and different causes might apply to different deficits.

Here, we examine low-level processes related to perceptual encoding. Behavioral studies that examined the quality of perceptual encoding in ADHD in the absence of attentional or executive involvement have found small and inconsistent differences (see [30] for a review). On the other hand, other investigations have found evidence for self-reported impairments in perceptual function in ADHD participants [31, 32], or in the general population with ADHD traits [33], as well as deficits in color processing and self-reported visual function in ADHD [34]. These findings are not necessarily contradictory, as perceptual deficits might emerge when attention or executive function is simultaneously taxed.

Therefore, we believe it is important to use a task that taxes both perceptual function and either attention and/or executive function, but that allows for a dissociation of the respective processes. This dissociation is difficult, as has been described in the study of autism [35]. In ADHD, there have been a few attempts to dissociate perceptual function from attention within a single task [36–38]. For example, [37] compared letter displays with or without distractors and found that ADHD participants had lower performance only when distractors were present. However, spatial covert attention was similar across ADHD and controls, leading the authors to suggest that perceptual interference or crowding is increased ADHD.

It is still unknown whether perceptual function is impaired when executive function is simultaneously taxed. A study by [39] used a face discrimination task where they probed perceptual noise by manipulating distractor saliency and probed top-down executive control by parametrically manipulating discrimination difficulty. In difficult discriminations, the reaction time difference between high-salience and low-salience distractors was comparable in children with ADHD to that in healthy children and adults; however, in easy discriminations, children with ADHD were slower to respond when presented with low-salience distractors. These results suggest similar perceptual interference due to distractor salience in ADHD and Controls, but a higher threshold in ADHD for activating executive control of attention. A problem with [39] is that face stimuli are high-dimensional and have content at many levels, complicating the separation between perceptual, attentional, and executive function. Another complication is that if the observer uses only 2 response keys in a task-switching paradigm, an error could be either due to a failure to switch or to a successful switch followed by a perceptual or attentional error [40].

Here, we attempted to characterize deficits in early processes of perceptual encoding in ADHD and dissociate them from executive deficits using a visuo-motor decision-making paradigm with task-switching which avoids the complications listed above. By using a total of 8 possible buttons out of which only 2 were relevant on a given trial, our response paradigm allowed for *task-irrelevant motor output* (TIMO), a new measure of executive control deficits. We defined a perceptual error as a press of the wrong button among the 2 relevant ones. We optimized the quantitative characterization of perceptual function by: a) using simple stimuli with feature dimensions orientation and color, thus minimizing high-level cognitive effects; b) varying stimuli parametrically along a continuum to estimate psychometric curve parameters (standard in perceptual psychophysics but still relatively rare in the study of ADHD [5, 36, 37, 39, 41, 42]); c) using an efficient stimulus selection method to minimize the number of trials needed for accurate estimation of parameters [43]. Broadly, our work follows a recent proposal to apply four levels of analysis to computational psychiatry: development of behavioral tasks, fitting of computational models, estimating parameters, and classification for diagnosis [44].

## Methods

### Approach

20 ADHD and 20 Control adult participants took part in our experiment. Stimuli were colored ellipses; each display contained one stimulus on the right of the fixation dot and one on the left. The participants performed yes-no discrimination (more precisely called yes-no classification or categorization) [45]. Specifically, the participants performed either fine orientation discrimination (was the cued ellipse clockwise or counterclockwise relative?) or fine color discrimination (was the cued ellipse more yellow or more blue?). The cue was 100% valid. In this task, participants had to rely on their internal memorized references, here for vertical and respectively the mid-level green in between yellow and blue.

We distinguish our design from other, perhaps more common, psychophysical tasks. For instance, 2AFC discrimination tasks could ask for a comparison between the 2 presented stimuli, for instance: ‘was the stimulus higher in contrast tilted to the right or to the left?’ [46]. We also distinguish our task also from orientation discrimination tasks in which, across trials, only 2 orientations are shown to be discriminated (say, −20 and +20), and other experimental manipulations are of interest.

Every trial started with a symbolic feature dimension cue, informing the participant which feature dimension was relevant on that trial. Simultaneously presented was a spatial cue (a line segment), informing the participant which side of the screen was relevant on that trial (Figure 1a). To better detect failures of spatial or feature switching, we used a response paradigm in which, on each trial, only 2 of 8 response keys were relevant, depending on the spatial and the feature cue; any other key press counted as task-irrelevant motor output (TIMO). Separately in each condition and for each participant, we used a Bayesian adaptive method to select maximally informative stimuli (see “Target stimulus generation”). This method allowed us to estimate the psychometric curve parameters with relatively few trials.

**Figure 1:**
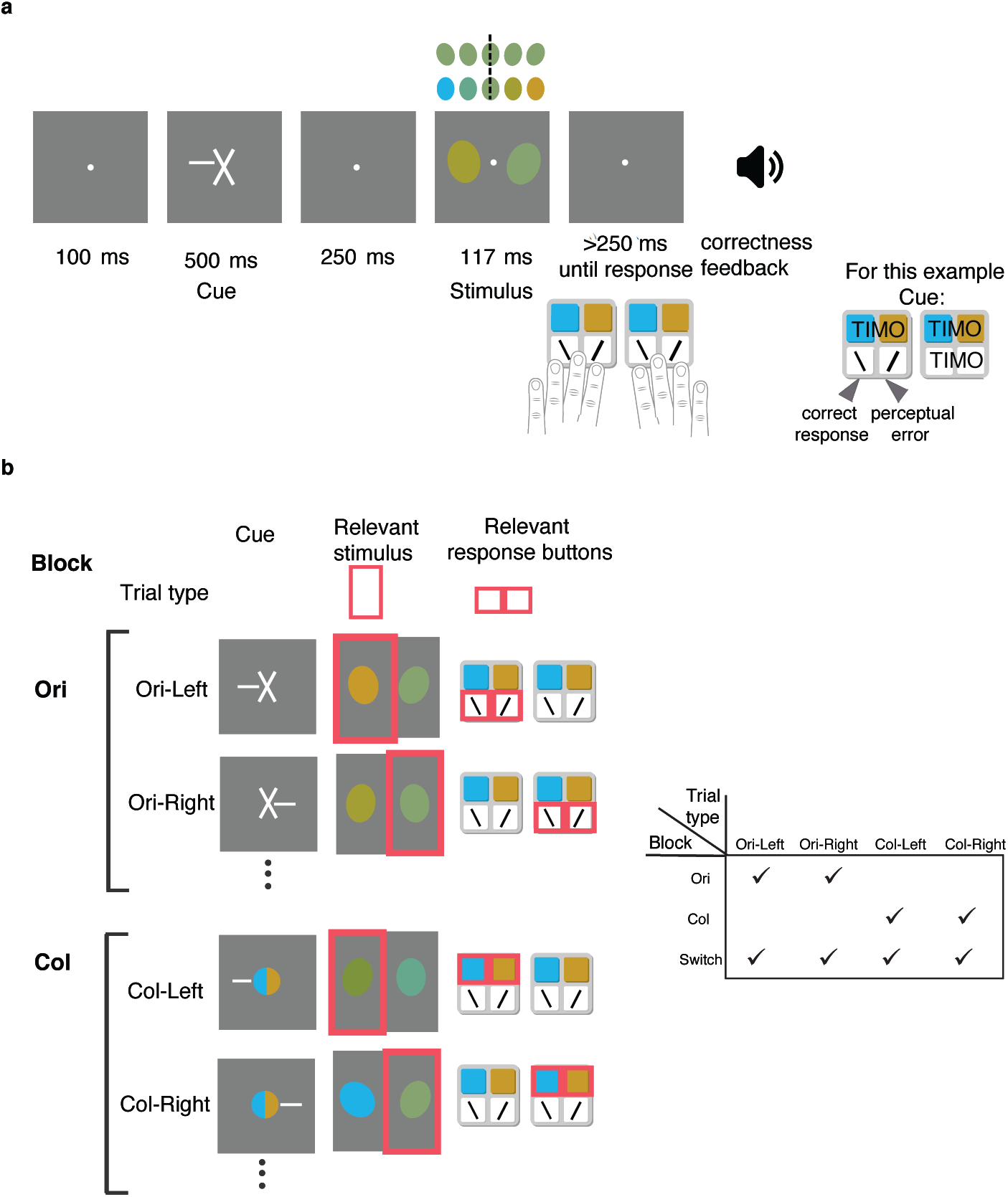
Task design. **(a)** Trial sequence example. A feature dimension cue indicated whether orientation (cross) - depicted here- or color (colored circle) was relevant, while a simultaneous endogenous spatial cue (line segment) indicated which side (left or right) was relevant. Thus, the participant received one of 4 possible cue screens. We always chose the spatial cue randomly. The participant had to respond whether the orientation of the ellipse on the relevant side was clockwise or counterclockwise with respect to vertical, or whether its color was more yellow or more blue, with the associated set of keys (left or right). The color and orientation continua are shown above the stimulus screen, with the dashed line at vertical and respectively mid-level green. To respond, the participant could press any one of 8 keys but only 2 were task-relevant on a given trial. The participant received correctness feedback. **(b)** (Left) Cue - relevant stimulus - relevant response buttons pairings for the 4 types of trials as they arise from the 4 feature and spatial cues combinations (2*2). Relevant is marked with pink for visualization only. Pressing any other button would result in TIMO. (Right) During Ori and Col blocks, only 2 types of trials are possible, while during Switch blocks all 4 trial types are possible.

Each participant experienced three types of blocks: Ori, Col and Switch. In Ori blocks, the feature dimension cue was always orientation. The spatial cue was randomly chosen on each trial, yielding 2 possible trial types: Ori-Left and Ori-Right (Figure 1b). We analyzed the Ori-Left and Ori-Right trials together as the Ori condition. In Col blocks, the feature dimension cue was always color and again the spatial cue was randomly chosen on each trial, yielding 2 possible trial types, Col-Left and Col-Right, which we grouped together for analysis into the Col condition. In Switch blocks, all 4 trial types were possible. We will refer to the orientation and color trials in switch blocks as the OriS and ColS conditions, respectively, and to the difference between no-switch and switch blocks as a difference in (executive) load.

An observer’s sequence of computations in the task can be conceptualized as a perceptual decision-making stage (stimulus encoding, affected by attention, and inference), followed by executive processing (rule retrieval and response execution) (Figure 2). The parametric variation of stimulus strength allowed us to estimate perceptual variability *σ* (or noise, the inverse of slope/sensitivity) as a main metric of perceptual function, and the 8-button response paradigm allows us to estimate task-irrelevant motor output as a main metric of executive function. In addition, we characterized behavior using other psychometric curve parameters, median reaction time, and reaction time variability.

**Figure 2:**
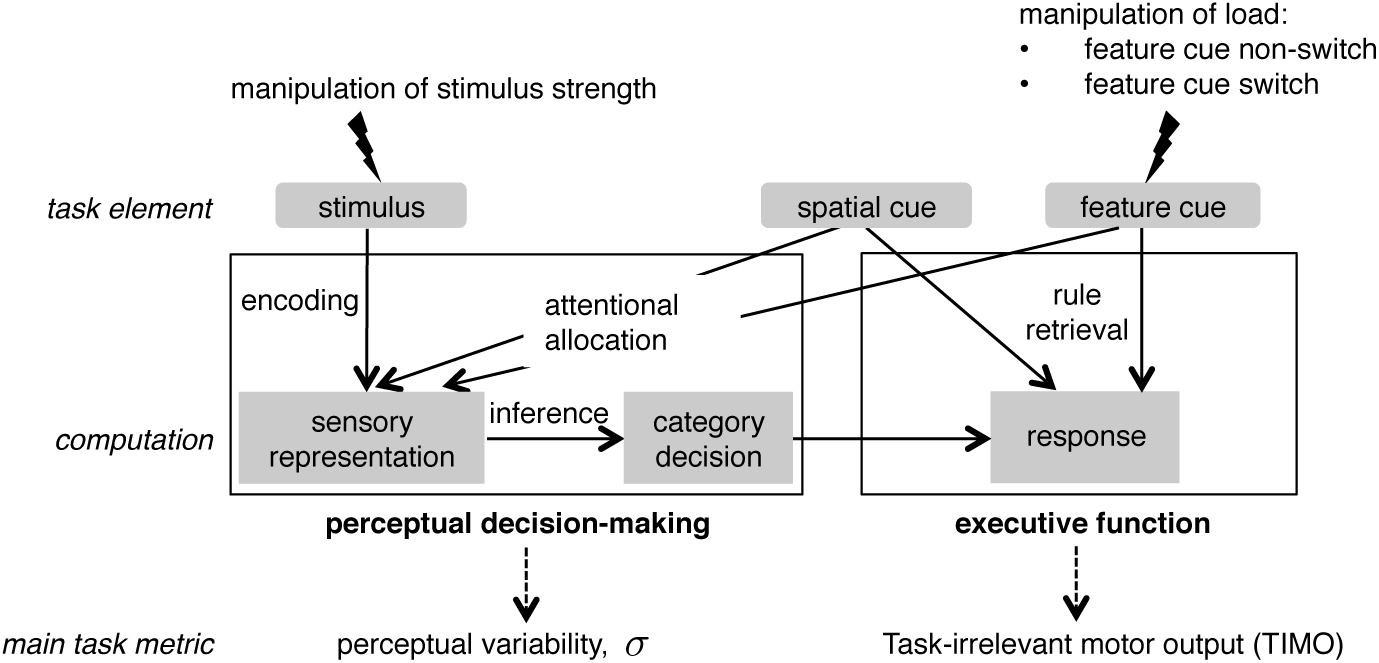
Dissociation of perceptual and executive processes. Schematic of the perceptual and executive processes that may play a role in this task, and the corresponding task metrics.

While usually a noise parameter (equivalent to our perceptual variability) in psychometric curves reflects a mix of sensory and decision noise [47], we believe that here the perceptual variability parameter for orientation and color is likely additionally modulated by attentional allocation. Previous studies showed modulation of psychometric curve parameters by attention, though either in different tasks such as target detection [48], 2AFC orientation discrimination [49], or color-change detection [50], or examined exogenous attention [51], or had other stimulus strength manipulation, such as contrast [52, 53] (for reviews see [54, 55]).

### Experiment

#### Participants

We recruited all participants through local advertisements, including flyers and newspaper and radio advertisements. Information on the participants is presented in Supplementary section “Demographic and clinical information”. Participants in both groups were matched as much as possible by age, sex, and education (see Table S1). 20 ADHD participants (12 female) of mean age 35.3 (SD: 10.0, range: 21 to 55) and 20 control participants (11 female) of mean age 32.5 (SD: 6.1, range: 19 to 44), with no statistical difference between their ages (Wilcoxon rank-sum test, *p* = 0.78), participated. 17 out of the 20 ADHD participants presented the combined subtype, and 3 the inattentive subtype. All participants spoke English and had normal or corrected-to-normal vision. We asked every participant before they started if they were colorblind. One participant was excluded because of color blindness. All participants provided informed consent. The study conformed to the Declaration of Helsinki and was approved by the Institutional Review Board of New York University School of Medicine.

#### Psychiatric assessment and diagnosis

None of the participants with ADHD were prescribed or took stimulant medication within 2 months of participating in the study. Participants with comorbid anxiety or unipolar depressive disorders were included as long as the symptoms at the time of evaluation were mild or in remission. Participants with bipolar disorders, psychotic disorders, substance use disorders, and neurologic disorders were excluded. For all adults, the diagnostic procedure included both clinician administered and self-administered scales. A trained clinician assessed every participant using the Adult ADHD Clinician Diagnostic Scale (ACDS) v.1.2, the Adult ADHD Investigator Symptom Rating Scale (AISRS), the Clinical Global Impressions-Severity of Illness (CGI-S) Scale, and the M.I.N.I International Neuropsychiatric Interview. All participants also completed the Adult ADHD Self-Report Scale (ASRS v.1.1.), the Adult ADHD Quality of Life (AAQoL) Scale, the World Health Organization Disability Assessment Schedule (WHODAS-II), and the Behavior Rating Inventory of Executive Function Adult Version (BRIEF-A). These scales have been extensively validated [56–59].

#### Apparatus

We displayed stimuli on a 23-inch (58.42 cm) Acer T232HL LCD monitor of resolution: 1920 × 1080 pixels and 60 Hz refresh rate (1 frame lasting 16.7 ms). We used a Kinesis Freestyle2 split keyboard. Participants used a head rest located at approximately 55 cm from the screen; this meant that 1 degree of visual angle (dva) subtended approximately 34 pixels. Stimulus presentation and response collection were controlled by a Windows computer running Matlab 7.1 (MathWorks, Massachusetts, USA) with Psychtoolbox3 [60–62] and EyeLink [63].

For 10 out of 20 ADHD participants and 10 out of 20 control participants, we monitored their fixation and recorded their eye movements. The rationale for not eye tracking all participants was a mixture of lack of sufficient time on the participants’ side and balanced design on the experimenters’ side. The eye tracker was calibrated using the five-point calibration routine before every block. We recorded eye movements using a remote infrared video-oculographic system (EyeLink 1000 Plus; SR Research, Ltd, Mississauga, Ontario, Canada) with a 1 kHz sampling rate. We set the heuristic filtering option OFF.

#### Stimuli

The background was mid-level gray (28.7 cd/m^2^). The stimuli were ellipses with area of 1600 pixels^2^ and 0.55° eccentricity, and thus with a major axis of 50 pixels and minor axis of 41 pixels. For the non-target ellipse, the orientation was randomly drawn from a von Mises distribution centered at 0 with *κ* = 30, and then divided by 2, approximately equivalent to a Gaussian distribution with mean 0 and a standard deviation of about 5°. The color of the non-target ellipse was based on a uniformly drawn sample that was used to linearly interpolate between blue and yellow in CIE L*a*b* (CIELAB) color space, with blue as [78 −30 −40], corresponding in RGB space to [0 167 255] and yellow as [78 0 80] corresponding in RGB to [200 130 0]. For each color, lightness was always kept constant at *L* = 78. Indeed, measured luminance was *∼* 39 cd/m^2^. The target stimulus was specified on a trial-to-trial basis using the Bayesian algorithm described below.

#### Target stimulus generation

The orientation and color of the target stimulus were based on the participant’s previous responses according to an adaptive procedure, a type of Bayesian staircase, applied separately for each of the 4 conditions. We used Acerbi’s Matlab implementation of the Ψ method called Psybayes [43], based on [64] with extentions to include the lapse rate [65]. This procedure maintains a posterior distribution over the parameters and updates it after each trial on which the participant pressed one of the 2 task-relevant buttons. The next stimulus value is chosen to minimize the entropy of the updated posterior given the stimulus, averaged over the participant’s possible responses weighted by their respective probabilities [64]. Each one of these 4 Bayesian staircases generated on every trial a unitless value *w* within the range [-0.5, 0.5] that was converted to stimulus values: to 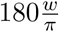 degrees in orientation trials and to [*L*, (*w* + 0.5)*a*_yellow_ + (0.5 *-w*)*a*_blue_, (*w* + 0.5)*b*_yellow_ + (0.5 *-w*)*b*_blue_] in color trials. Thus, target stimulus values fell within an orientation range of −30 deg to 30 deg and within a blue to yellow range from [78 −30 −40] to [78 0 80]. We defined the space of parameters that Psybayes constructs the posterior on: for *µ*, we used a linear grid of 51 points from −0.5 to 0.5, for *σ*, a logarithmic grid of 25 points from 0.002 to 0.8 and for *λ* a linear grid of 25 points from 0 to 0.3.

#### Trial sequence

(Figure 1a). A trial sequence started with the simultaneous appearance of a feature dimension cue and a spatial cue, presented for 500 ms. The feature dimension cue for orientation consisted of 2 white line segments, each of length approximately 1 dva, crossing at the center, with orientations tilted ±26.6° with respect to vertical; for color, it consisted of 2 semi-circles (divided vertically, right one yellow, left one blue) joined to form a circle of radius approximately 0.3 dva. Simultaneously, a spatial cue was presented, which consisted of a horizontal line segment of length approximately 0.5 dva emanating from the center of fixation to the left or to the right. We chose 500 ms to ensure sufficient time for the deployment of endogenous feature-based attention [66]. Following a delay of 250 ms consisting of the presentation of a central fixation circle of radius 0.12 dva, 2 ellipses appeared at 2.5 dva to the right and left of a central fixation circle. The stimuli were presented on the screen for 117 ms, followed by another delay period of 250 ms.

After the post-stimulus delay, the participant had to respond about the target ellipse via a specific key press out of a total of 8 keys (Figure 1a). On any given trial, 6 of these 8 keys are irrelevant. For orientation, the participants were instructed to press one of the 2 labeled keys for clockwise (CW) or counterclockwise (CCW), using the left keypad for the left spatial cue and the right one for the right spatial cue. For color, they had to press one of 2 labeled keys to indicate whether the ellipse was more yellow or more blue, also using the left or respectively right keypad depending on the spatial cue. Figure 1b shows all these 4 possible cue-response mappings. The direction of the spatial cue was randomly drawn on each trial, so participants used their right hand approximately half the time. After the response, auditory feedback was provided for 200 ms: a 1200 Hz tone if the participant had pressed the correct key, and a 500 Hz tone if the participant had pressed any of the 7 incorrect keys.

#### Training

Before they began the experiment, participants were guided step by step through the different parts of instructions. The experimenter read the instructions on the screen (presented in Figure S1a) out loud. To remind subjects of the stimulus-response pairings, a sheet with these pairings was posted on the wall of the psychophysics room (Figure S1b). In total, participants performed 40 training trials: a short orientation only block (’O’) of 10 trials, a short color only block of 10 trials and a short switch block (’S’) of 20 trials. The experimenter was present with the participants during the training to observe responses, provide further feedback and answer questions. Participants repeated the set of all 40 training trials until they achieved a performance greater than 65%.

#### Experiment structure

Afterwards, they performed 8 blocks of about 100 trials each in the order ‘O-C-S-S-S-S-C-O’ or ‘C-O-S-S-S-S-O-C’, with 30 seconds breaks in between blocks. Changes in block type were signaled with a screen with the instruction ‘In this block, your job is to report ORIENTATION’ for O blocks, or ‘In this block, your job is to report COLOR’, for C, or ‘In this block, your job is to report either ORIENTATION or COLOR’, for S, with each feature dimension word followed by its associated symbol. In total, participants completed 800 non-aborted trials, approximately 200 in each one of the four conditions, Ori, Col, OriS and ColS (from S blocks).

### Statistical analyses

For most metrics, we report median values and 95% bootstrapped confidence intervals. Across 50000 iterations, we took samples with replacement from and of the same size as the original data with Matlab’s `randsample` and calculated the median of each of those sets of samples. The the 2.5th and 97.5th quantiles of the distribution of medians across iterations were taken as the 95% confidence intervals.

#### Three-way mixed-design ANOVA

To determine the differences between groups and the 2 experimental conditions of load and feature, we used three-way mixed-design ANOVA with two repeated measures, since we have one “between - participants” variable (group) and two “within - participants” factors (feature - Ori vs Col and load-No-switch vs Switch). Beforehand, we log transformed the measures that were lower bounded by 0. When we assumed shared parameters between No-switch and Switch and thus we had only one “within - participants” factor, we used two-way mixed-design ANOVA. We implemented the ANOVAs in SPSS with “General linear model: repeated measures”. For post-hoc comparisons, we adjust the significance level according to the Sidak correction to 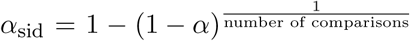. For the three-way mixed-design ANOVA, we performed, unless otherwise specified, 12 planned pairwise comparisons in Matlab: Wilcoxon rank-sum tests between groups (one for each condition, 4 in total), and Wilcoxon signed-rank tests for conditions within a group (4 per group, 8 in total). We used the Sidak correction for multiple comparisons, decreasing the significance level to *α* = 0.0043 for post hoc comparisons following the three-way mixed-design ANOVA or respectively *α* = 0.0127 following the two-way mixed-design ANOVA.

#### Pairwise correlations

To correct for multiple comparisons when examining the pairwise correlation matrix of the performance measures, we used a method from Nyholt et al. [67]. If *M* is the total number of measures, the number of effective comparisons will be decreased more if the measures are more highly correlated, as captured in a higher variance of the eigenvalues *λ*_obs_ of the correlation matrix, which we calculated with Matlab’s function `eig`. Then, 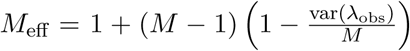. As in [67], *M*_eff_ is used in the Sidak correction (a slightly less conservative alternative to the Bonferroni correction), modifying the significance level to 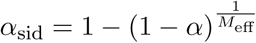.

#### Linear regression

We implemented multivariate linear regression with Matlab’s `fitlm`.

#### Logistic regression for classification

We fit the logistic regression coefficients with Matlab’s `glmfit` with input ‘binomial’ and the link parameter ‘logit’. For a given participant, we used the task metrics and the fitted coefficients with `glmval` to get *p*(Diagnosis), which was then thresholded at 0.5 to predict the 0 or 1 ADHD diagnosis.

#### Stratified 10-fold cross-validation

In order to assess the use of this logistic regression classifier for out-of-sample prediction, we calculated the cross-validated accuracy. We did stratified 10-fold cross-validation, in which each fold had 4 participants, 2 ADHD and 2 Controls; thus, we trained the classifier to find the coefficients over 36 participants and tested over 4 and calculated the mean accuracy across folds. We did 1000 runs of this stratified 10-fold cross-validation to allow for different random assignments of participants into folds and took the mean accuracy over runs.

### Parameter fitting

#### Psychometric curves and parameters

We fitted psychometric curves to trials on which a participant pressed one of the 2 relevant buttons. *s* denotes the normalized stimulus value on a given trial (ranging between [-0.5, 0.5]). We use the following form of the psychometric curve [68]:

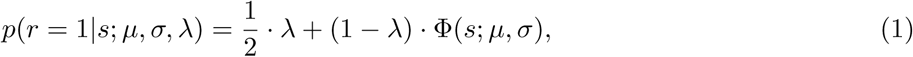

where *r* = 1 stands for a response “clockwise” (orientation) or for “more yellow” (color). The parameters are the point of subjective equality (PSE or bias), *µ*, the inverse slope (or noise) parameter, *σ* - which both are inputs to the Gaussian cumulative density function (Φ) - and the lapse rate, *λ*. We had 4 conditions, Ori, OriS, Col and ColS and thus 4 psychometric curves.

#### Parameter estimation and model choice

We performed maximum-likelihood estimation of the psychometric curve parameters *µ, σ*, and *λ*. The likelihood of a parameter combination is the probability of the data given that parameter combination; we denote the log likelihood by LL. We assumed that trials are independent of each other and thus we summed the log likelihoods across all trials. We fitted orientation and color trials separately; thus the following log likelihoods apply to either set of trials. In the main model, we assumed that *µ* and *λ* are shared across both load conditions (No-switch and Switch), whereas *σ* might differ. These assumptions had both a practical and a principled motivation. Assuming that parameters are shared between conditions reduced the number of parameters to 8 and made parameter estimates more reliable. Moreover, if *µ* reflects an overall bias and *λ* a generic lapsing process, we did not expect them to change with load. For a model without these assumptions, and a model comparison, see Supplementary section “Further information on psychometric curves and parameters”. The log likelihood for trials in a given feature dimensions becomes

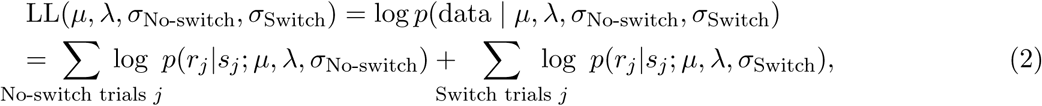

where *s*_*j*_ and *r*_*j*_ are the stimulus and the participant’s response on the *j*th trial, respectively. To estimate the parameters, we searched on a grid with 201 values in each dimension: for *µ* linearly spaced from −0.2 to 0.2, for *λ* logarithmically spaced from 0.0001 to 0.3, and for each *σ* logarithmically spaced from 0.002 and 0.5.

#### Reaction times

For fitting ex-Gaussian distributions to reaction times, we used a custom made script modeled after an existent software package [69].

#### Data and code availability

Clinical data is not available beyond diagnosis labels, experiment code is available upon request and behavioral data and analysis code are available at https://github.com/lianaan/Perc_Var.

## Results

We attempted to dissociate perceptual from executive deficits in ADHD with a new visuo-motor decision-making task with a task-switching component. This task yielded two main measures: task-irrelevant motor output and perceptual variability.

### Task-irrelevant motor output (TIMO)

TIMO refers to the trials when participants pressed one of the 6 irrelevant keys and hence such responses most likely reflect a failure of proper stimulus-response rule retrieval. TIMO was quite low overall (mean ± sem: 0.06 ± 0.01); ADHD participants produced a higher proportion of TIMO (0.079± 0.018) relative to Controls (0.041± 0.008). Figure 3a presents a breakdown of TIMO by condition. A three-way mixed-design ANOVA on log TIMO with between-participants variable group and within-participants factors load (No-switch and Switch) and feature (Ori and Col) reveals a significant effect of group 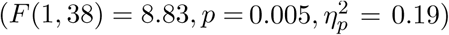, a significant effect of load 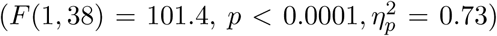, and no significant effect of feature 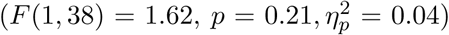. Neither of the two-way interactions nor the three-way interaction were significant (*p* > 0.06). In particular, the group × load interaction was not significant 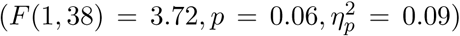; thus, we did not find that switching between feature dimensions carries a higher cost in ADHD. Next, we performed 12 post-hoc planned comparisons: within each group, Wilcoxon signed-rank tests for Ori versus OriS, Col versus ColS, Ori versus Col, and OriS versus ColS, and between groups, Wilcoxon rank-sum tests for Ori, OriS, Col, and ColS. After Sidak correction (*α* = 0.0043), no between-group comparisons were significant (*p* > 0.0046). The within-group load comparisons were all significant (*p* < 0.002). No within-group feature comparisons were significant (*p* > 0.07). Taken together, these results validate TIMO as a metric of interest for executive control.

**Figure 3:**
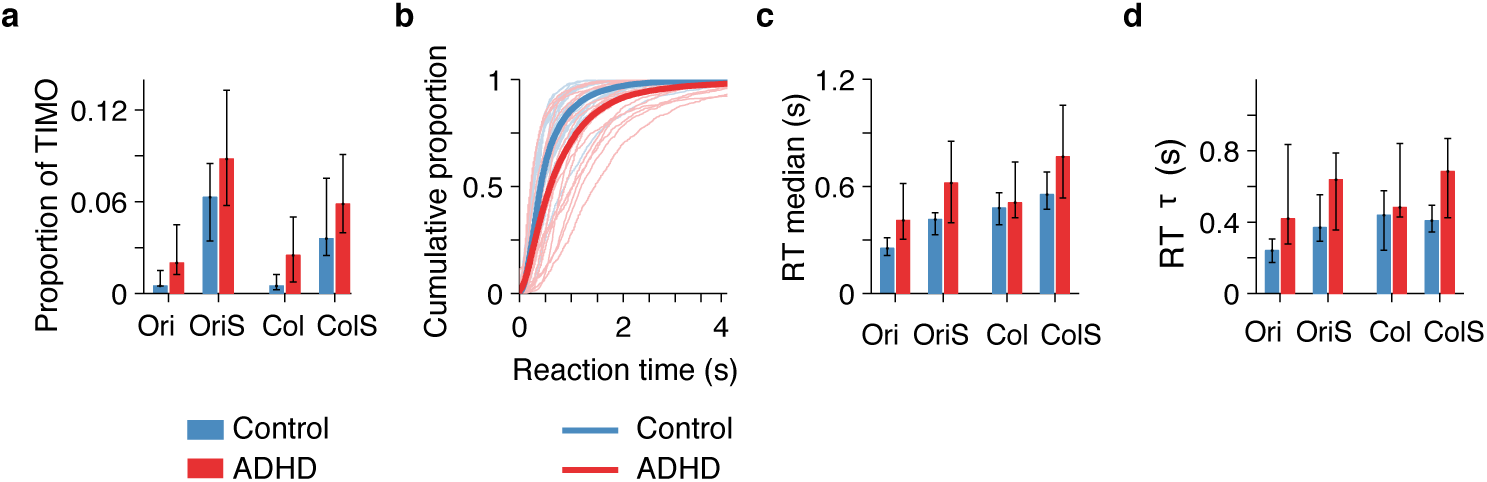
ADHD participants had higher TIMO and longer and more variable reaction times. **(a)** Proportion of TIMO across conditions. Here and elsewhere, values represent medians across participants and error bars the bootstrapped 95% confidence intervals. **(b)** Empirical cumulative density functions of reaction times, collapsed across all conditions. Thin lines: individual participants. Thick lines: median for the RT distribution collapsed across all participants in a group. **(c)** Reaction time median by condition and group. Throughout the paper, we use RT median because reaction time distributions are not Gaussian. **(d)** Reaction time variability metric, the *τ* parameter from ex-Gaussian distribution fits, by condition and group.

In the OriS and ColS conditions, the majority of TIMO seemed to be feature errors (Figure S2b). Relative to the instructions on a given trial, the 6 irrelevant keys subdivide into 2 that represent spatial errors, 2 feature errors and 2 that represent both spatial and feature errors. We did not delve into these distinctions since overall TIMO was quite low.

### Reaction times

ADHD participants showed longer reaction times (RTs; Figure 3b). Three-way mixed-design ANOVA on log RTs revealed a significant effect of group 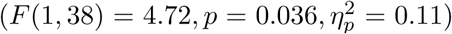, and significant effects of load 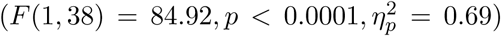 and feature 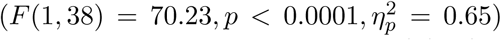. In addition, we found significant group × feature 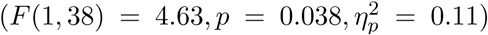 and load × feature interactions 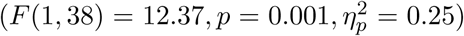, but not a significant group × load interaction 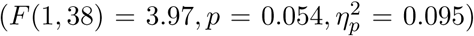. After Sidak correction (*α* = 0.0043), none of the between-group comparisons were significant (*p* > 0.019). The effects of within-group load and feature on log RTs were all significant both within Control and within ADHD (*p* < 0.001). Higher RTs for Col than Ori could be due to the fact that the Ori responses are intuitively mapped to left and right, while the Col responses are arbitrarily mapped as blue to left and yellow to right.

Higher RT variability (or intra-individual variability) in ADHD has been found consistently [9] and has been generally thought to reflect cognitive processes separate from higher median RTs [9, 70] (but see [71] for an opposing account). The term RT variability has been used to refer to different aspects of RT distributions [9]; here, we fitted ex-Gaussian distributions [72] and used the *τ* parameter as a measure of RT variability. The *τ* parameter has been shown to capture the tendency of ADHD participants to have a higher proportion of abnormally slow responses [9, 70, 72]. Before committing to the ex-Gaussian, we verified that it captures the data better than the log-normal and gamma distributions (see Supplementary section “Further information on reaction times”). Three-way mixed-design ANOVA on log *τ* revealed a significant effect of group 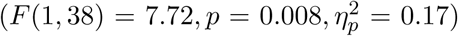, an effect of load 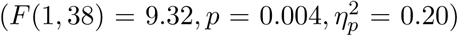 and an effect of feature 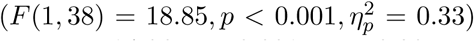. The only significant interaction was between load and feature 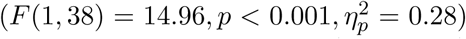. After Sidak correction (*α* = 0.0043), none of the between-group comparisons were significant (*p* > 0.006). Within Controls, the effects of load and feature on log RT *τ* were significant for Ori vs. OriS and Ori vs. Col (*p* < 0.001). Within ADHD, no effects of load or feature were significant (*p* > 0.02). We confirmed the pattern of higher RT variability in ADHD with a non-parametric measure, RT iqr (see Supplementary section “Further information on reaction times”).

Overall, we found that ADHD participants had longer and more variable reaction times, consistent with previous work [9, 72, 73]. However, RT-related differences across groups are usually difficult to interpret because they might encompass multiple processes, including sensory encoding, decision time, speed-accuracy trade-offs, stimulus-response rule retrieval, response preparation, and response execution (unless some of these processes are disentangled with drift diffusion models [71, 74]). The effect of load on RT and RT *τ* does seem to suggest that on Switch trials, more time is spent on executive processes, here stimulus-response rule retrieval, response preparation, and response execution, relative to No-Switch trials.

### Psychometric curve parameters

We confined the following analysis to the trials in which participants pressed one of the 2 relevant keys. Because of the Bayesian stimulus selection method, each participant received a different set of stimuli for each condition (see Figure S6) and thus proportion correct is largely stable across conditions and participants (mean ± sem: 0.811 ± 0.007, Figure S2A) and thus not informative. Instead, we fitted a psychometric curve to non-TIMO trials in each condition [75]. Thus, the parameters of the psychometric curves captured the differences in performance across conditions and participants. The normalized orientation and color continua spanned the interval [-0.5, 0.5]. Each psychometric curve has three parameters: a point of subjective equality *µ*, perceptual variability *σ*, and a lapse rate *λ* (Figure 4b,c and Figure S8d). Non-zero *µ* represents a tendency to choose one option more than the other. The parameter *σ* is a composite of sensory noise and decision noise, and also reflects the quality of the allocation of spatial attention, and of feature attention in switch blocks. Higher *σ* denotes a reduced ability to discriminate between small variations within a feature. The parameter *λ* reflects trials with lapses in attention or erroneous motor output. In our main model, we assumed that *µ* and *λ* are independent of load; we confirmed this assumption by comparing to a model without these assumptions (“full” model) in the Supplementary section “Further information on psychometric curves and parameters”. The parameters *σ* and *λ* might trade off against each other, although this is less of a concern in our main model than in the “full” model.

**Figure 4:**
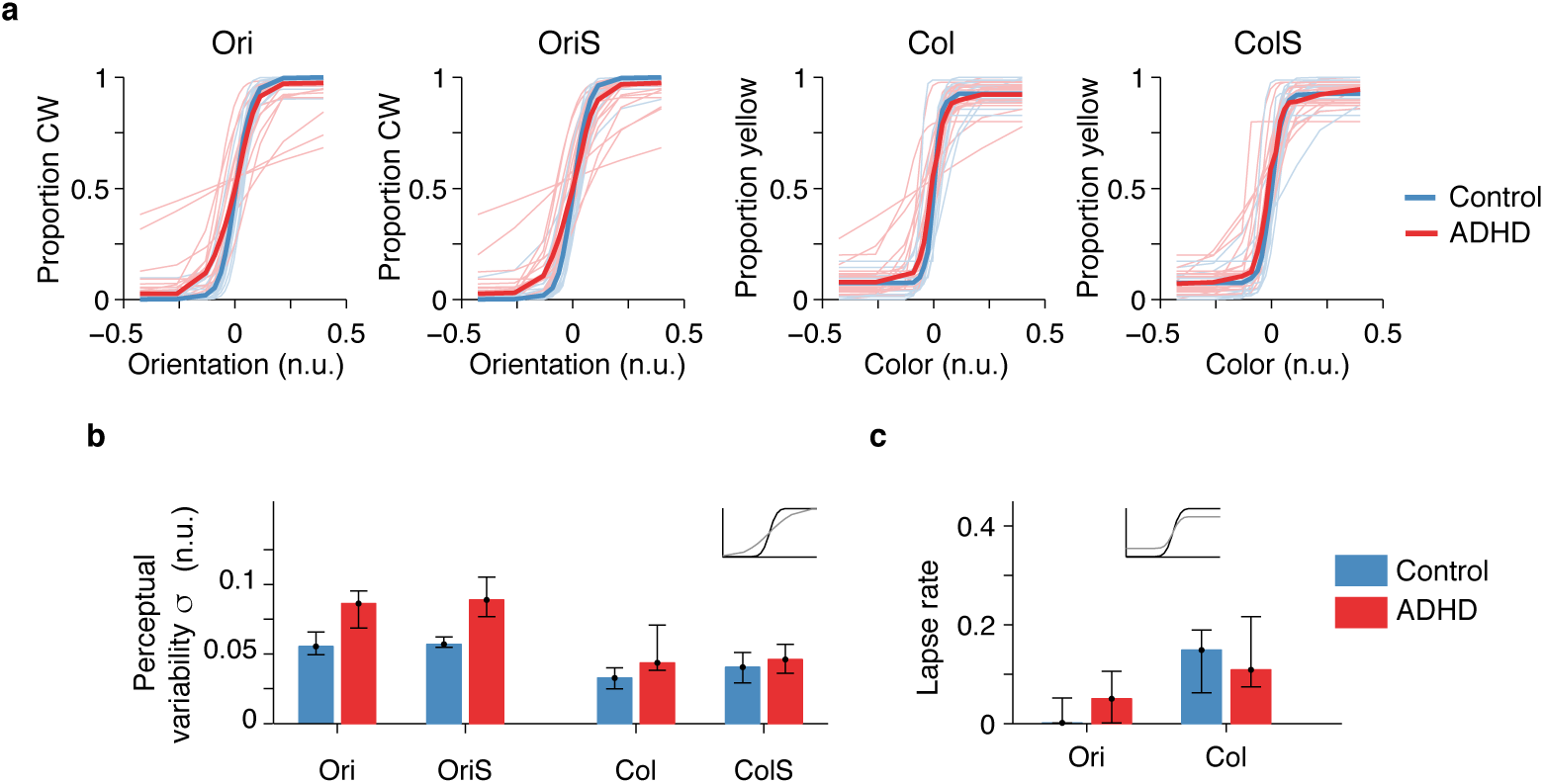
Fitted psychometric curves and parameters. ADHD participants had higher perceptual variability. **(a)** Psychometric curve fits across all conditions. Here and elsewhere, n.u. stands for normalized units. Thin lines: individual participants. Thick lines: medians for each group. For fits overlaid on top of data, see Figure S8. **(b)** Perceptual variability parameter values, medians and bootstrapped 95% confidence intervals. Top inset plot: black psychometric curve has low noise, while the grey has higher noise. **(c)** Lapse rate. Top inset plot: black psychometric curve has low lapse, while the grey has higher lapse.

Three-way mixed-design ANOVA on log *σ* showed a significant effect of group 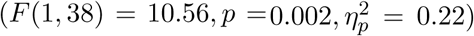, a significant effect of feature 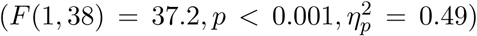, but not of load 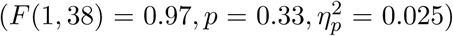. The effect of group × load was not significant 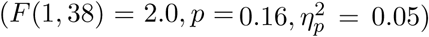. Because the normalization to the (arbitrary) stimulus range is specific to each feature dimension, the values of *σ* cannot be meaningfully compared between orientation and color: a different stimulus range would have changed the *σ* values without changing the observer’s true perceptual variability. Therefore, only the within-feature post-hoc comparisons are meaningful, giving a corrected significance level of *α* = 0.0065. Then, the between-group comparisons were significant for both Ori and OriS (*p* < 0.0005), but not for Col (*p* = 0.0083) or ColS (*p* = 0.28). No post-hoc comparisons with load were significant neither within Controls nor within ADHD (*p* > 0.01). Higher *σ* for orientation in ADHD could result from worse low-level sensory encoding (e.g. higher neural noise), lower covert endogenous attention, higher decision noise, or even noise in the inference process about the perceptual category. The lapse rate reflects responses that are independent of the stimulus, such as lapses of attention, but could also trade off with the *σ* parameter. Two-way mixed-design ANOVA on log *λ* showed a large effect of feature 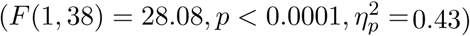, but no significant effect of group 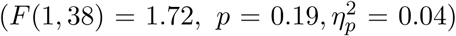 and no significant group × feature interaction. After Sidak correction (*α* = 0.0127), we found that Control (*p* < 0.0001) and ADHD (*p* = 0.011) participants tended to lapse more for color than for orientation, possibly because the stimulus-response mapping is less intuitive. Results for *µ* are in the Supplementary section “Further information on psychometric curves and parameters”. In conclusion, the parametric variation of low-level stimulus variables combined with stimulus optimization revealed robust perceptual deficits in ADHD, especially for orientation.

A possible cause of the increased perceptual variability in ADHD could be that ADHD participants were slower to learn the task. To check for learning, we fitted two psychometric curves for each condition, one to the first half of the trials and one to the second half. The *σ* parameters across participants, conditions and time are presented in Figure S9. Visually, we notice a slight improvement in perceptual variability in the second half of the trials (Figure S9). To quantify it, we performed a 4-way mixed-design ANOVA on log *σ* with time as an additional factor. We found an effect of time 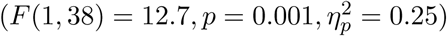. We do not find a significant group × time interaction 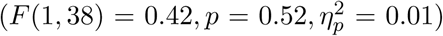 and thus we have no evidence for a differential learning pattern for ADHD relative to Controls.

### Correlations across metrics

Next, we asked whether behavioral metrics are correlated with each other (Table 1). For this analysis, we collapsed across groups; per participant, we averaged each behavioral metric across all four conditions. We found that the perceptual variability parameter *σ* is significantly correlated with TIMO, RT median, and RT *τ*, with high effect sizes. Note that the perceptual variability parameter and TIMO were computed from different sets of trials, therefore reducing the probability that their correlation is spurious. In addition, a breakdown of some of these correlations by group, symptom type, and condition is presented in Supplementary section “Breakdown of correlations”.

**Table 1:** Pairwise Spearman correlations across log task metrics (both behavioral and clinical). Both TIMO and perceptual variability are significantly correlated with several other variables. Boldfaced denotes significance after multiple-comparisons correction (*α* = 0.0089, see Methods).

| | TIMO | RT | RT $\tau$ | Perceptual<br>variability ($\sigma$) | Lapse rate ($\lambda$) | GEC |
| --- | --- | --- | --- | --- | --- | --- |
| TIMO |  |  |  |  |  |  |
| RT | **$\rho = 0.46$**<br>$p = 0.003$ | | | | | |
| RT $\tau$ | **$\rho = 0.42$**<br>$p = 0.007$ | **$\rho = 0.84$**<br>$p < 0.0001$ | | | | |
| Perceptual<br>variability ($\sigma$) | **$\rho = 0.41$**<br>$p = 0.0085$ | **$\rho = 0.55$**<br>$p = 0.0003$ | **$\rho = 0.57$**<br>$p = 0.0002$ | | | |
| Lapse rate ($\lambda$) | **$\rho = 0.46$**<br>$p = 0.003$ | $\rho = 0.23$<br>$p = 0.15$ | $\rho = 0.17$<br>$p = 0.30$ | $\rho = 0.28$<br>$p = 0.08$ | | |
| GEC | **$\rho = 0.53$**<br>$p = 0.0005$ | $\rho = 0.25$<br>$p = 0.12$ | $\rho = 0.34$<br>$p = 0.03$ | **$\rho = 0.50$**<br>$p = 0.0009$ | $\rho = 0.30$<br>$p = 0.06$ | |
| ACDS | $\rho = 0.40$<br>$p = 0.01$ | $\rho = 0.31$<br>$p = 0.05$ | **$\rho = 0.45$**<br>$p = 0.004$ | **$\rho = 0.51$**<br>$p = 0.0008$ | $\rho = 0.18$<br>$p = 0.26$ | **$\rho = 0.80$**<br>$p < 0.0001$ |

So far, we have characterized behavioral differences between ADHD and Controls in our task. Next, we asked if behavioral metrics relate to common clinical ones, namely the General Executive Composite score (GEC), as assessed by the Brief-A questionnaire (self-reported, [76]), as well as the (ACDS) scores (clinician interview, [56]). The GEC and ACDS (Table S2) are meant to be continuous measures of executive control and symptom severity, respectively. Both GEC and ACDS revealed strong correlations with TIMO, suggesting that TIMO could serve as a behavioral marker of executive deficits. GEC and ACDS were both also strongly correlated with perceptual variability. In addition, ACDS (but not GEC) was correlated with RT *τ*, which provides a graded counterpart of the robust finding of increased RT variability in ADHD [9]. A linear regression of GEC as a function of behavioral metrics (*R*^2^ = 0.38) showed only TIMO as statistically significant (Table S7a), reinforcing our interpretation of TIMO as reflecting failures of executive function. A linear regression of ACDS as a function of the same metrics (*R*^2^ = 0.33) only showed significance for RT *τ* (Table S7b), suggesting that GEC and ACDS, despite being strongly correlated (Figure S11), could capture distinct aspects of impairment [77]. However, the determinant of the correlation matrix of these measures is 0.11, nearing 0 and thus signaling multicollinearity [78]. Therefore, we have to be cautious in interpreting the individual contributions of these regressors. Nevertheless, these results suggest that our behavioral metrics capture to some extent the same processes as clinical metrics, while having the advantage of avoiding the potential subjectivity inherent in questionnaires.

### Classification of participants

Finally, we asked how accurately we can classify a given participant as ADHD or Control based purely on behavioral task metrics. Figure 5 depicts these results. A logistic regression using only the perceptual variability parameter yielded a classification accuracy of 77.5%, with a hit rate (sensitivity) of 75% and a false-alarm rate (1 minus specificity) of 20% (Figure 5a). A logistic regression classifier based on TIMO only had an accuracy of 62.5%, with a hit rate of 65% and a false-alarm rate of 40%; using both perceptual variability and TIMO improved the accuracy to 82.5%, with a hit rate of 80% and a false-alarm rate of 15% (Figure 5a). Of note, while these variables are correlated, the determinant of their correlation matrix is 0.82, far enough from 0 that collinearity should not be a problem [78]. Adding more regressors (RT, RT *τ* and lapse) did not yield further improvement (80.0%); this is not surprising in light of multicollinearity. Hence, we consider perceptual variability and TIMO as the main regressors of interest. In order to assess the use of this logistic classification for out-of-sample prediction, we did stratified 10-fold cross-validated and found mean accuracies of 77.1% with perceptual variability as the only regressor, 63.1% with TIMO only, 77.8% with both perceptual variability and TIMO and 70.0% with all metrics. The relatively high classification performance suggests that our task has potential as a diagnostic tool.

**Figure 5:**
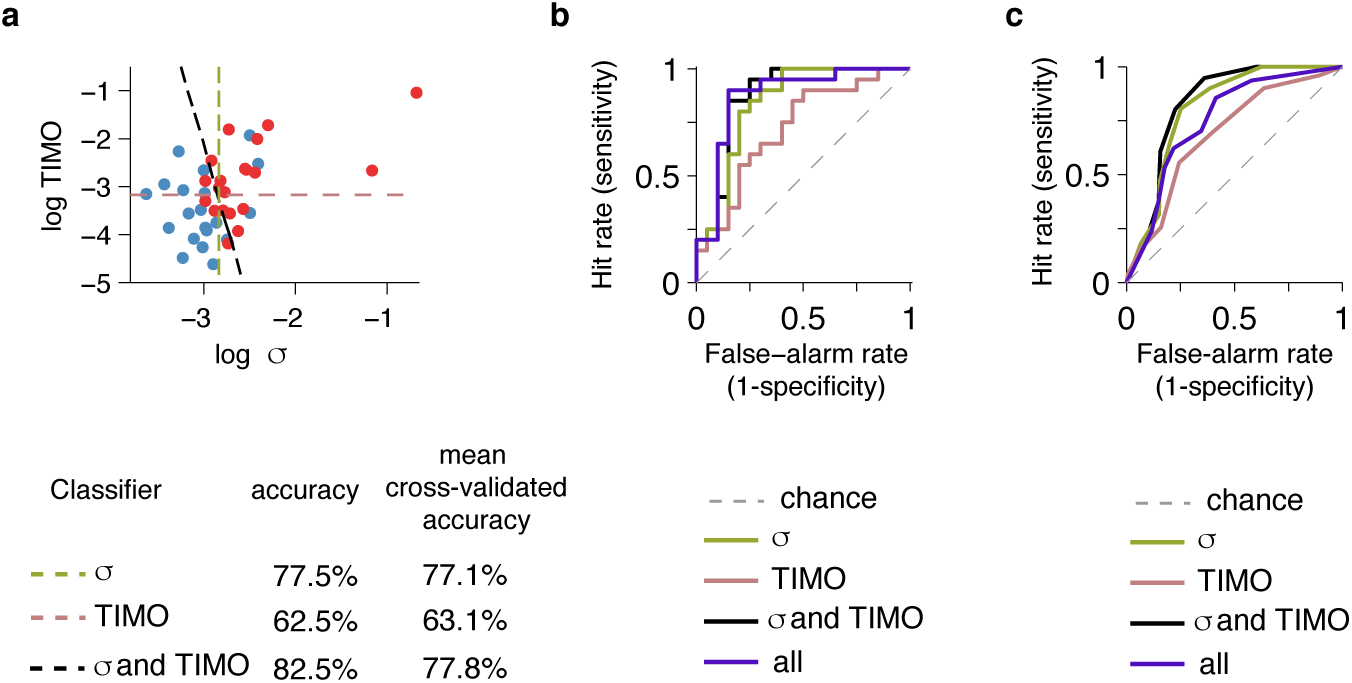
Logistic regression based on task metrics can classify participants into ADHD and Controls with accuracies larger than 70%. **(a)** Dots: combinations of log TIMO and log perceptual variability (*σ*) across participants. Dashed lines: logistic regression classifiers trained on log *σ* only (olive), TIMO only (old rose) and both (black). **(b)** Full ROC curves obtained by varying the diagnosis threshold for the three classifiers in **(a)**, as well as for one based on all 5 behavioral metrics (purple). **(c)** Full ROC curves, this time with stratified 10-fold cross-validation, for the same classifiers as in **(b)**.

In addition to thresholding at 0.5 to get diagnosis and subsequently accuracy as above, we also thresholded *p*(Diagnosis) at linearly spaced values between 0 and 1 and plotted the resulting receiver operating characteristic (ROC) curves, both without (Figure 5b) and with stratified 10-fold cross-validation (Figure 5c). As expected, the ROC curve for the classifier all metrics shows the best performance (highest area under the curve (AUC)) without cross-validation, but its performance degrades for out-of-sample predictions in the cross-validated case.

## Discussion

In this study, we dissociated stimulus encoding (perceptual) from response selection (executive) deficits in ADHD with a new visuo-motor decision-making task with a task-switching component. To better separate executive deficits from perceptual and attentional failures, we used 8 response keys, 6 of which were irrelevant on any given trial (TIMO). To assess perceptual precision, we used simple stimuli that varied continuously along one dimension. We used and an adaptive stimulus selection method [43] that reduced the number of trials needed for accurate parameter estimation (relative to, for instance, the method of constant stimuli); reducing the number of trials is crucial when running the ADHD population. We found differences between ADHD and Controls in our main task metrics, TIMO (Figure 3a) and perceptual variability (Figure 4b), as well as median reaction times and reaction time variability (Figure 3c and d). We found correlations of these behavioral metrics with clinical metrics (Table 1) and were able to classify participants into ADHD and Controls with high accuracy solely on the basis of our main behavioral metrics (Figure 5).

Our finding of higher TIMO in ADHD could be due to more spatial switching errors or more feature switching errors, but it is hard to quantify these contributions since TIMO was overall relatively low. It is conceivable that a less intuitive stimulus-response mapping for orientation distrimination (stimulus oriented towards left/respond with key on the left), or types of stimuli that require spatial integration [79–82] or cross-modal switching [83], or more complex forms of task switching [6, 7], would produce larger differences on a TIMO-like executive function measure.

In line with previous work [9, 72, 73], we found that ADHD participants had longer and more variable reaction times. While accuracy was maintained to be approximately stable in all participants, perceptual variability was higher in ADHD, and thus the increased reaction times are not reflective of speed-accuracy trade-offs. In addition, our paradigm allowed us to analyze correlations across individuals between reaction time metrics and other metrics. The correlation between the perceptual variability parameter *σ* and median reaction time is consistent with a drift-diffusion model, in which slower accumulation of evidence simultaneously leads to lower accuracy and longer reaction times. Indeed, many studies have found slower drift rates in ADHD [24, 71, 74, 84, 85].

We found higher *σ* in ADHD than in controls. This parameter - which we called the perceptual variability parameter could be affected both by sensory encoding (affected by attention) and decision processes. Could the differences in *σ* be attributed to either type of process? Sensory and decision noise are usually confounded in the parameters derived from behavior in common discrimination tasks [47]. However, tasks exist in which the influences of sensory and decision noise can be separated [86, 87]. Additionally, neural data with high temporal resolution such as EEG or MEG could separate perceptual from decision-related variability as early vs late activity relative to stimulus onset [20, 88]. Decision noise in perceptual decision-making might be related to decision noise on action selection in reinforcement learning models of high-level cognitive tasks. [89] found increased decision noise (temperature parameter) in ADHD and later proposed this to underlie behavioral variability found in ADHD more generally [90]. Our result of increased perceptual variability parameter in ADHD is consistent with this general proposal, and extends it to include the possibility of an even lower-level correlate of behavioral variability.

Earlier studies examining perceptual function in isolation did not find differences between ADHD and Controls (see [30] for a review). Our result of higher perceptual variability in the ADHD group suggests that the encoding of visual stimuli is less precise than in Controls, at least when experimental conditions simultaneously tax other processes. In our case, participants had to allocate either spatial attention or both spatial and feature-based attention, as well as employ executive function by maintaining and acting on either 2 (no-switch) or 4 (switch) stimulus-response rules. Earlier studies examining covert spatial attention while attempting to minimize executive load did not find differences between ADHD and Controls [2–5]. While perceptual precision and attention might be comparable between ADHD and Controls when studied in isolation, it is possible that asking ADHD participants to simultaneously devote brain resources to other processes might allow for differences in perceptual variability to emerge.

### Possible lower-level neural correlates of behavioral variability in ADHD

Our results could speak to the question of low-level perceptual components interacting with measured executive control deficits, as we found a significant correlation between the perceptual variability parameter and the executive control metric TIMO. In particular, our results raise the possibility of a shared neural source of perceptual and executive function deficits, such as a lower signal-to-noise ratio in early brain areas. While ideas of lowered signal-to-noise ratio implemented through impaired dopamine and noradrenaline signaling in ADHD have been put forward before, they have been mainly confined to cerebellar, striatal and prefrontal regions [90–92]. Beyond that, one study found higher neural noise in the visual and auditory cortex of ADHD participants [20]. ADHD participants could have higher perceptual variability in orientation by having less selective orientation tuning of cells in V1; this was the mechanism proposed to underlie decreased orientation discrimination with aging in monkeys [93]. The list of regions with lower signal-to-noise ratio in ADHD could include deeper brain structures with roles in selecting relevant sensory stimuli and maintaining stimulus-response rule representations such as the thalamus [94–98], or even lower regions with roles in orienting of attention and behavioral flexibility, such as the superior colliculus [99, 100] or the locus coeruleus [101, 102]. However, these regions do not just modulate cortical representations but also receive substantial top-down inputs, so the source of the reduced signal-to-noise ratio could originate from either lower or higher-level brain regions.

Based on our data, we cannot establish whether the proposed low-level level correlate of behavioral variability is reflective of a diffuse deficit, of frontal-based executive function, or of impairments in endogenous attention reliant on fronto-parietal circuits. Nevertheless, our results make the case that low-level perceptual function in ADHD deserves further investigation and that future task designs can easily include assessments of perceptual function in conjunction with attention and executive function. Using simple rather than high-level cognitive stimuli has the advantage that they can be used in parallel human and animal studies. Studies on mouse [103, 104] and rat [105] models of ADHD will provide further insight into the neural circuits implicated in ADHD and how medications can alter these circuits [106, 107].

### Perceptual variability as a candidate diagnosis marker for ADHD

ADHD diagnosis still relies predominantly on self and sometimes collateral reports and widely accepted “psychomarkers” (also called “neurocognitive endophenotypes”) and biomarkers are lacking [108]. For our findings to have implications for clinical practice, it is necessary that our task metrics are predictive of clinical metrics. We found that this was indeed the case. First, based on perceptual variability alone, we were able to classify participants into ADHD and Control with cross-validated mean accuracy of 77.0% (including TIMO, 77.7%). Beyond binary classification, we also found strong correlations between behavioral metrics (*σ*, TIMO, and RT *τ*) and clinical ones (GEC and ACDS). Based on these correlations, the behavioral metrics in our task could be considered candidate psychomarkers for ADHD, similar to the performance on the CPT [109], response variability [8, 110], and drift rate [111], and along with potential biomarkers such as saccade patterns [112], microsaccade rate in specific tasks [33, 113, 114], pupil size [115], eye vergence [116] and cortical thickness [117]. While there is a long pipeline from task and metric to clinically useful assay [118, 119], simple behavioral paradigms and modeling applied to ADHD and other disorders could in the long term help refine diagnostic categories and inform and quantify the efficacy of treatment, as is the goal in computational psychiatry more broadly [44, 120, 121].

## Acknowledgements

This work was supported by NIH grant R01EY020958 to W.J.M. and an NYU SoM Applied Research Support Fund (ARSF) grant number 53101. We thank Terry Leon, Michael Silverstein, Saima Milli, Jonathan Yuh for assistance with protocol preparation, participant recruitment and clinical data management and analysis. We thank Ili Ma and two anonymous reviewers for comments on earlier versions of this manuscript. In addition, we thank Eero Simoncelli, Marisa Carrasco, Roozbeh Kiani, Mariel Roberts, and members of the Ma lab, especially Maija Honig, Luigi Acerbi, Will Adler, Bas van Opheusden, and Aspen Yoo, for useful conversations.

## Supplementary section

### S1 Demographic and clinical information

**Table S1:**
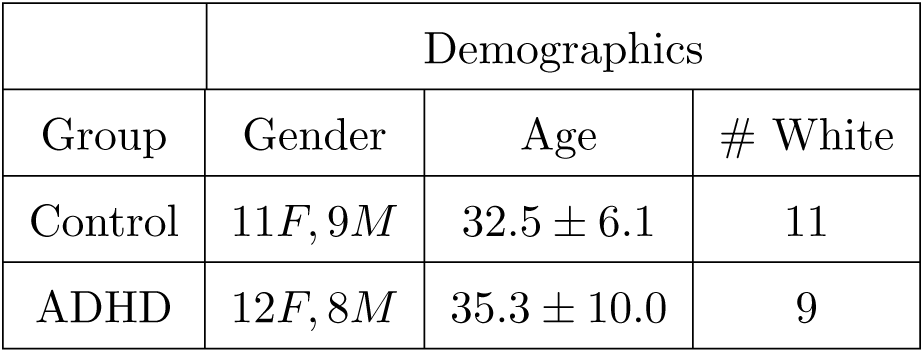
Demographic information of participants. Values represent mean and standard deviation.

**Table S2:**
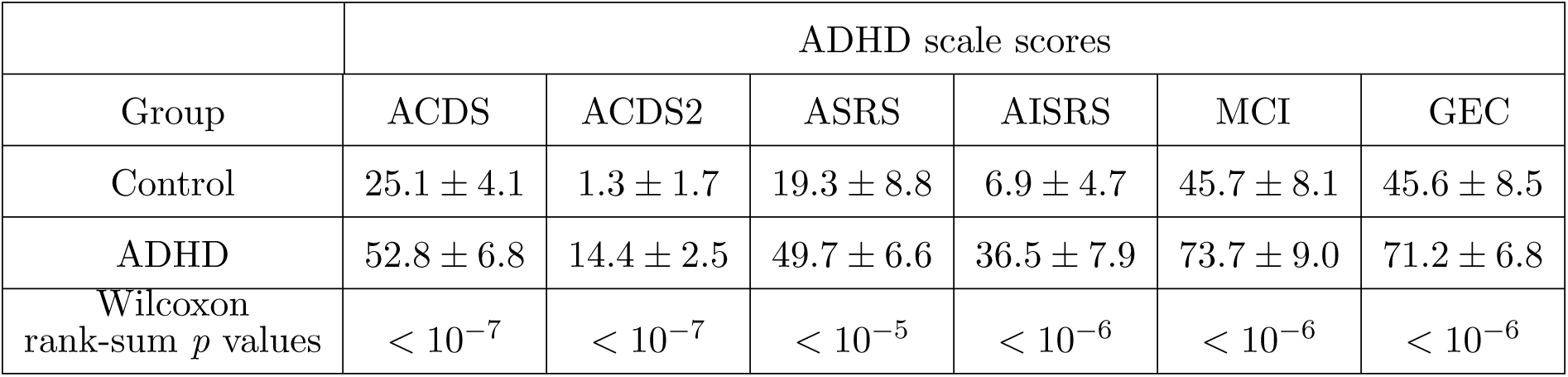
Psychiatric scores of participants. Values represent mean and standard deviation. ACDS denotes ACDS B1-B18, and ACDS2 B22-B39.

**Table S3:**
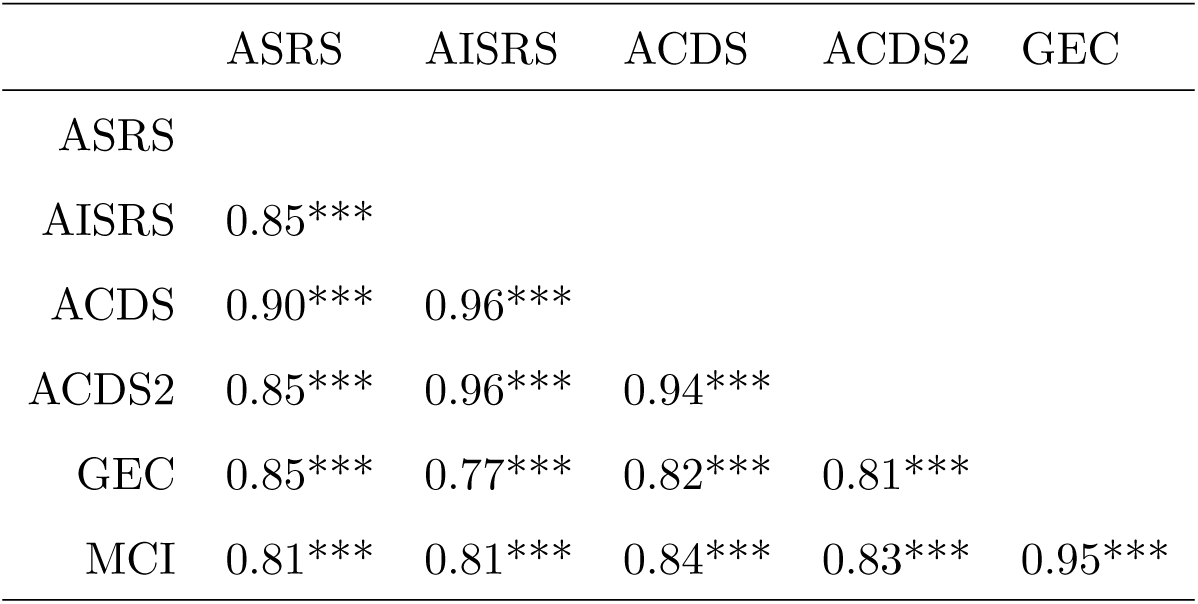
Spearman correlations across the scores for all diagnosis scales. 10 participants were excluded from this table because not all records were available. However, ACDS, ACDS2, GEC and MCI were available for all participants.

**Figure S1:**
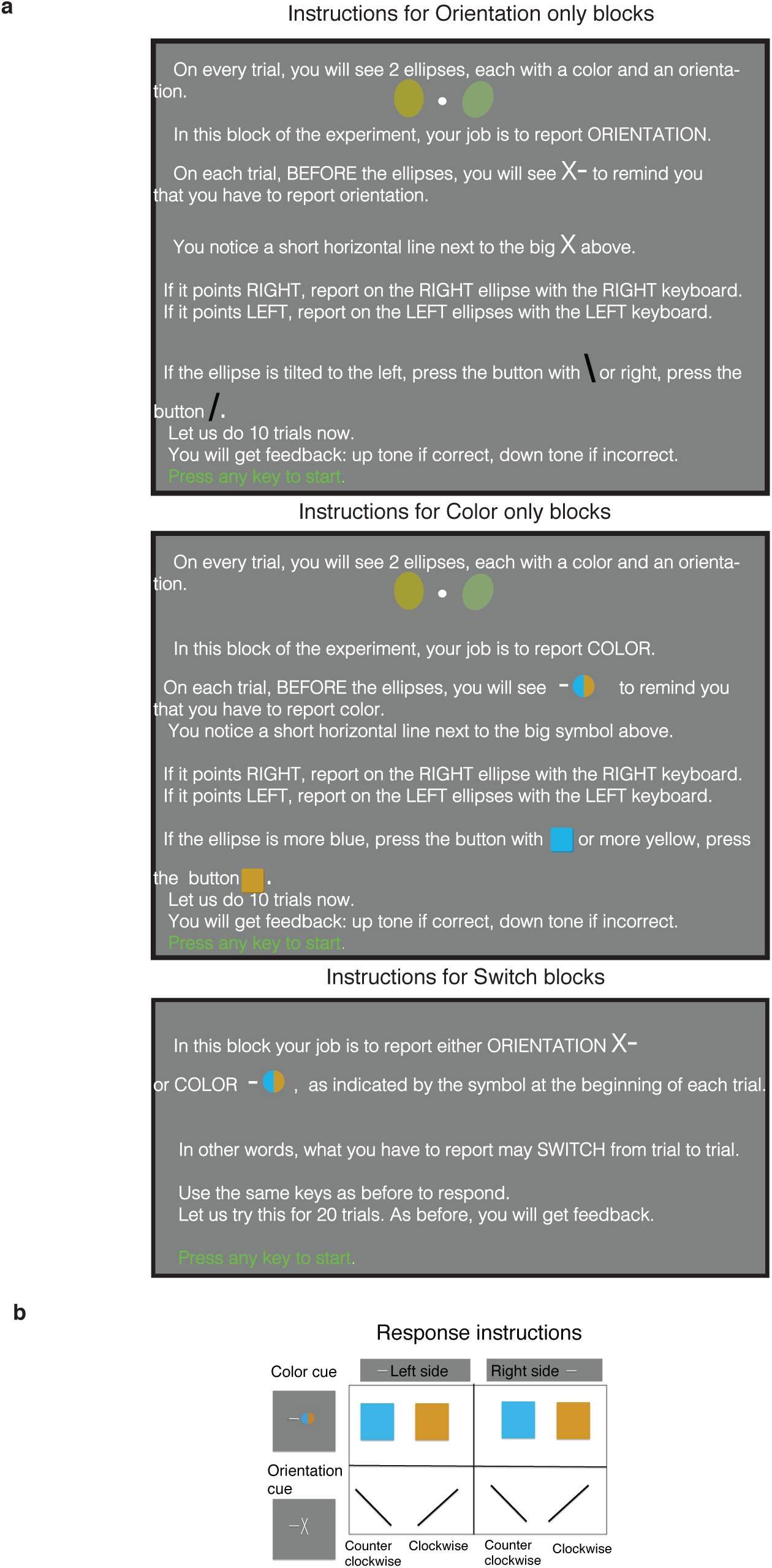
Training information. **(a)** Training instructions for the 3 different types of blocks: Orientation only, Color only and Switch. **(b)** Reminder of the stimulus-response pairings. A sheet containing this information was present on the wall of the psychophysics room within participants’ sight.

### S2 Further characterization of responses

Accuracy was maintained approximately constant across participants and conditions (mean ± sem: 0.811 ± 0.007) due to the Psybayes method of adaptive thresholding (Figure S2a). We further characterized the TIMO responses, first with a breakdown by error type, available for 32 participants (Figure S2b) and then according to the type of the previous trial (Figure S2c). In the Switch trials, the majority of TIMO seem to be feature errors, possibly because the mapping from left/right visual field to left/right keyboard was more intuitive than from feature dimension to top/down of a keyboard (Figure S2b). Lastly, we saw no clear pattern from breaking down proportion of TIMO by the type of the previous trial (Figure S2c).

**Figure S2:**
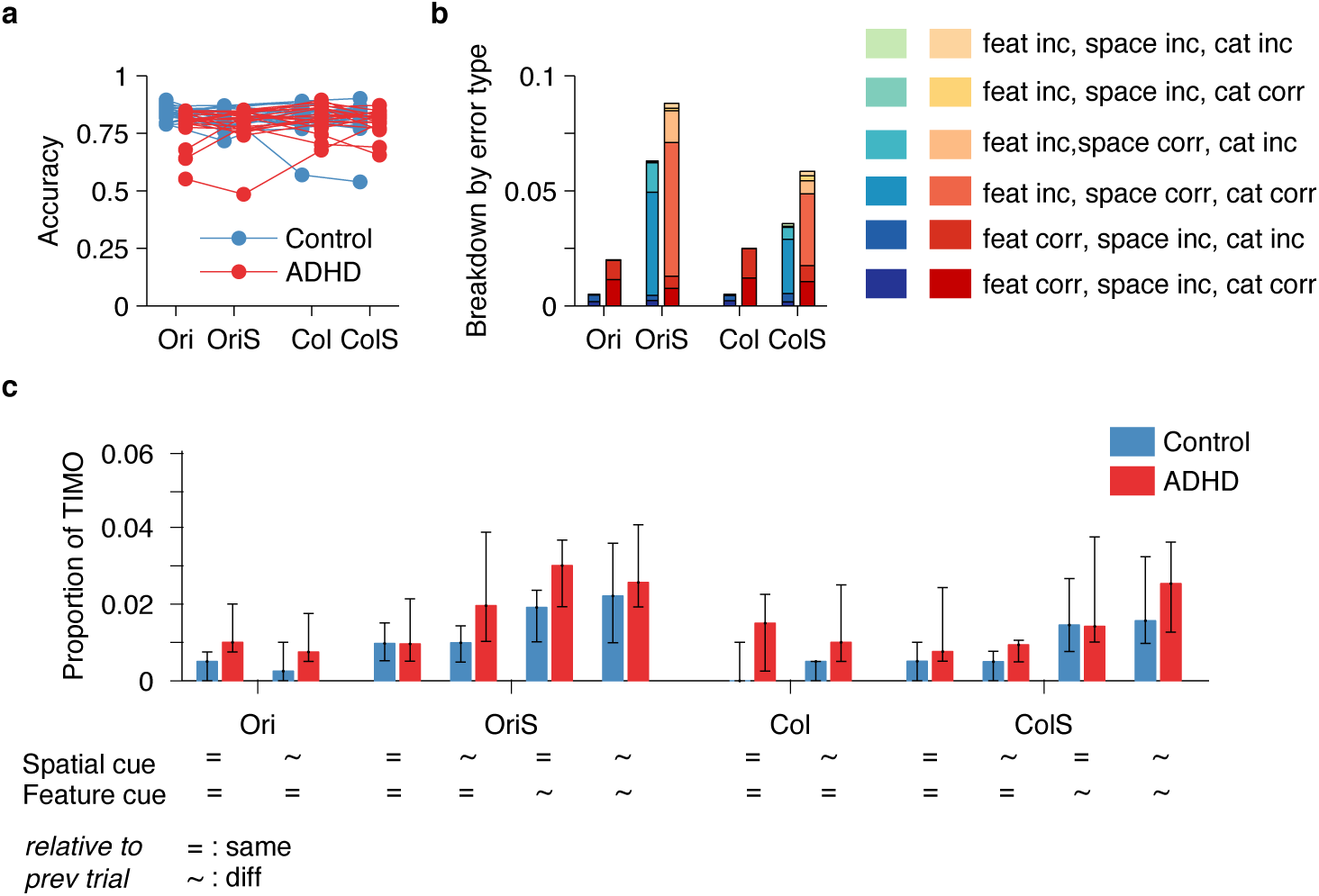
Further characterization of responses. **(a)** Accuracy as proportion correct on the trials when participants selected one of the 2 relevant keys. **(b)** The task irrelevant motor output from Figure 1, broken down by error type. For every condition, the first bar is Controls, and the second one ADHD. **(c)** Same, broken down by the type of the previous trial.

### S3 Further information on reaction times

#### S3.1 ex-Gaussian model

ex-Gaussian distributions are commonly fitted to reaction time data and are defined by adding 2 random variables, a Gaussian with parameters *µ* and *σ* and an exponential with parameter *τ*. While in our data *τ* showed an effect of group, neither log *µ*_RT_ 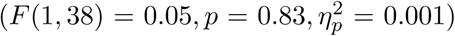 nor log *σ*_RT_ 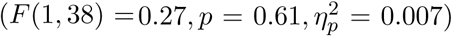 did, consistent with other studies that showed significant effects of group on *τ* but not on *µ*_RT_ (see [9] for a meta-analysis).

**Figure S3:**
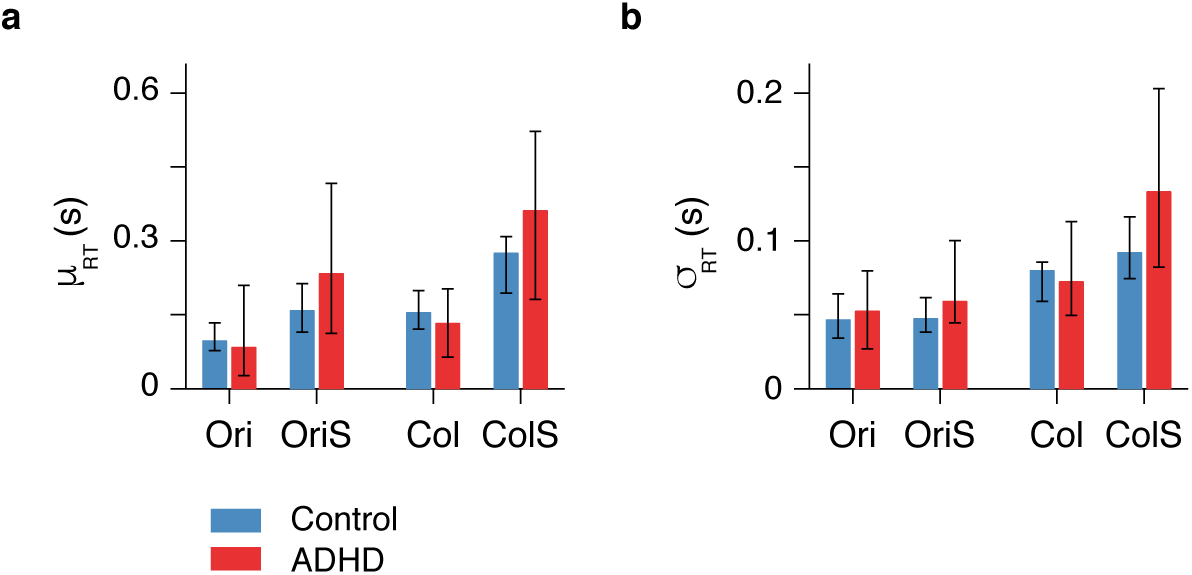
ex-Gaussian parameters fitted to the reaction time distributions across conditions and groups. **(a)** Gaussian mean *µ*_RT_, **(b)** Gaussian standard deviation *σ*_RT_, both for each task condition.

#### S3.2 Alternative models

While ex-Gaussian distributions are routinely used to fit reaction times, they are rarely compared to alternative distributions. We used the corrected Akaike Information Criterion (AICc) and the Bayesian Information criterion (BIC) to compare the ex-Gaussian fits with the fits of 2 other distributions on the positive real line: log-Normal and Gamma. These metrics are defined as 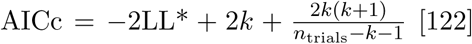 [122] and BIC = −2LL* + *k* log *n*_trials_ [123], respectively, where LL* is the maximum log likelihood, *k* is the number of free parameters, and *n*_trials_ is the number of trials. We found that indeed the ex-Gaussian distribution was a better fit than both the log-Normal (in median by 611 according to AICc and by 607 according to BIC) and the Gamma distribution (in median by 50 according to AICc and by 45 according to BIC); see Figure S4 for individual subjects.

#### S3.3 Non-parametric measure of RT variability

We complemented the results about RT *τ* (Figure 3) with a non-parametric robust measure of intraindividual reaction time variability, the reaction time inter-quartile range (iqr) (Figure S5). Three-way mixed-design ANOVA on log RT iqr’s revealed a significant effect of group 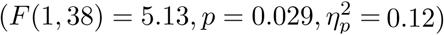, load 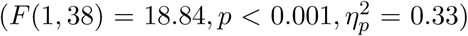, feature 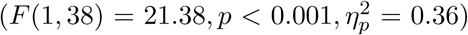, and a significant load × feature interaction 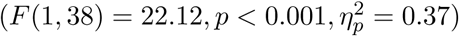. No other two-way interaction nor the three-way interaction were significant (*p* > 0.36). After Sidak correction (*α* = 0.0043), none of the between-groups comparisons were significant. Within Controls, the effects of load and feature on log RT iqr were significant for Ori vs OriS (*p* = 0.0015) and Ori vs Col (*p* < 0.001); within ADHD, the only significant effect was of load for Ori vs OriS (*p* = 0.002).

**Figure S4:**
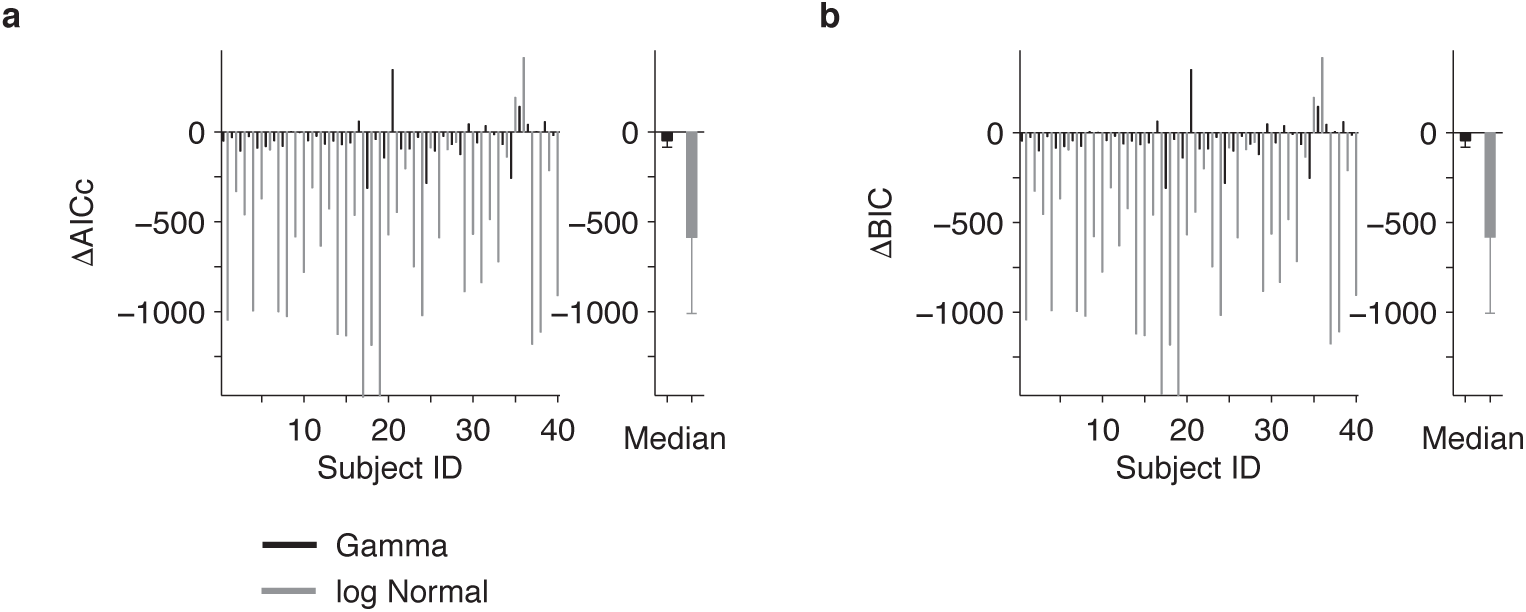
Model comparison justifies the parametrization of reaction times with the ex-Gaussian distribution. **(a)** The ex-Gaussian model has the lowest AICc across the population **(Right)** and for almost all individual subjects **(Left)**. **(b)** Same result for BIC.

**Figure S5:**
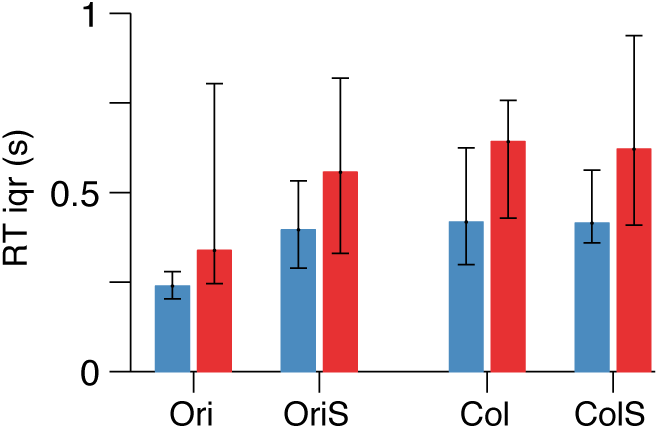
Reaction time variability is higher in ADHD also according to a non-parametric metric, RT iqr.

### S4 Further information on psychometric curves and parameters

#### S4.1 Stimuli sets

Figure S6 depicts the histograms of selected stimuli for each condition and each participant, optimized with the Bayesian stimulus selection method. As a consequence of this method, proportion correct is largely stable across conditions and participants (see Results), and the differences between participants were quantified through the psychometric curve parameters. In line with ADHD participants having higher perceptual variability, we see here that the collapsed histograms across all participants within a group show that Controls received a higher proportion of more difficult stimuli (higher bump around 0).

**Figure S6:**
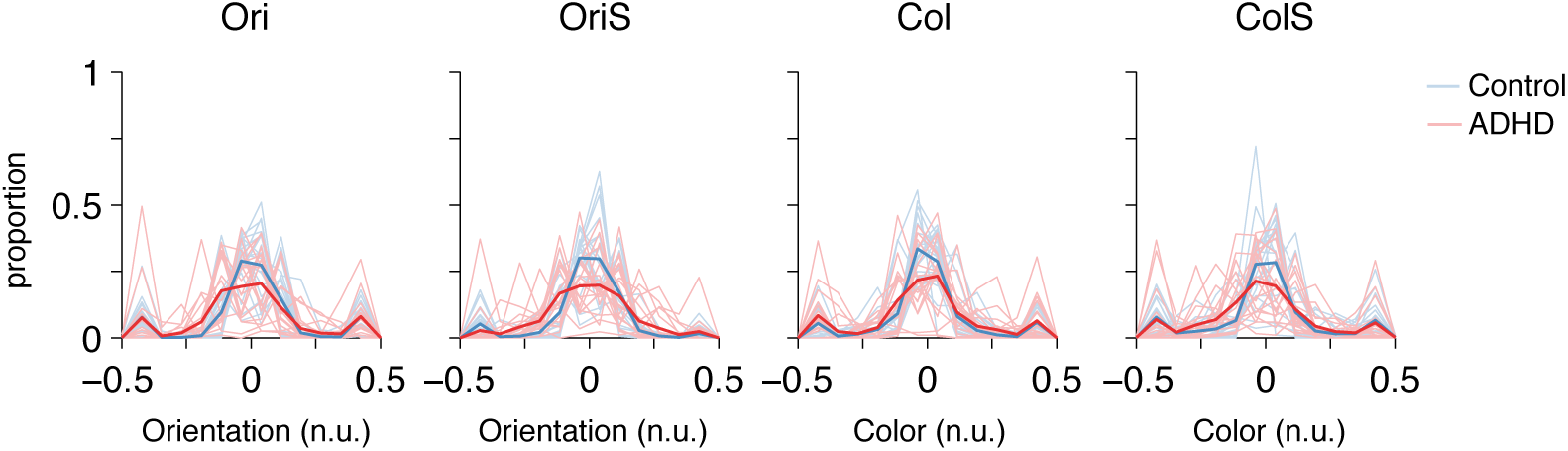
Distributions of stimuli across conditions and participants. Thin lines: individual participants. Thick lines: proportion of stimuli collapsed across all participants within a group.

#### S4.2 PSE

Figure S8d shows the estimates of *µ* (PSE) in the “shared” (main) model. Two-way mixed-design ANOVA on *µ* with within-group factor feature showed an effect of group 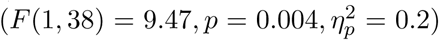, but no significant effect of feature 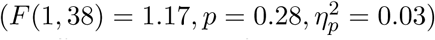 and not a significant interaction. After Sidak correction (*α* = 0.0253), no effects were significant. We chose to interpolate color values between blue and yellow since the S-cone pathway is of special interest in ADHD [124]. While we found an overall group effect on *µ*, after Sidak correction the post hoc effect for color failed to reach significance, thus making our results at this point inconclusive about whether ADHD participants have different S-cone dependent color processing.

#### S4.3 “Full” model, 12 parameters

While in the main or “shared” model with 8 parameters (Figure 4) we assumed that *µ* and *λ* were shared within a feature across load conditions, in the “full” model we did not constrain any parameters, yielding 12 parameters total.

As expected, the “full” model captured the data at least as well as the “shared” model. However, the “shared” model provided either a comparable (in median better by −1.5 according to AICc) or better (in median by −20 according to BIC) than the “full” model (Figure S7). This confirmed the plausibility of the shared-parameters assumption in the main model.

**Figure S7:**
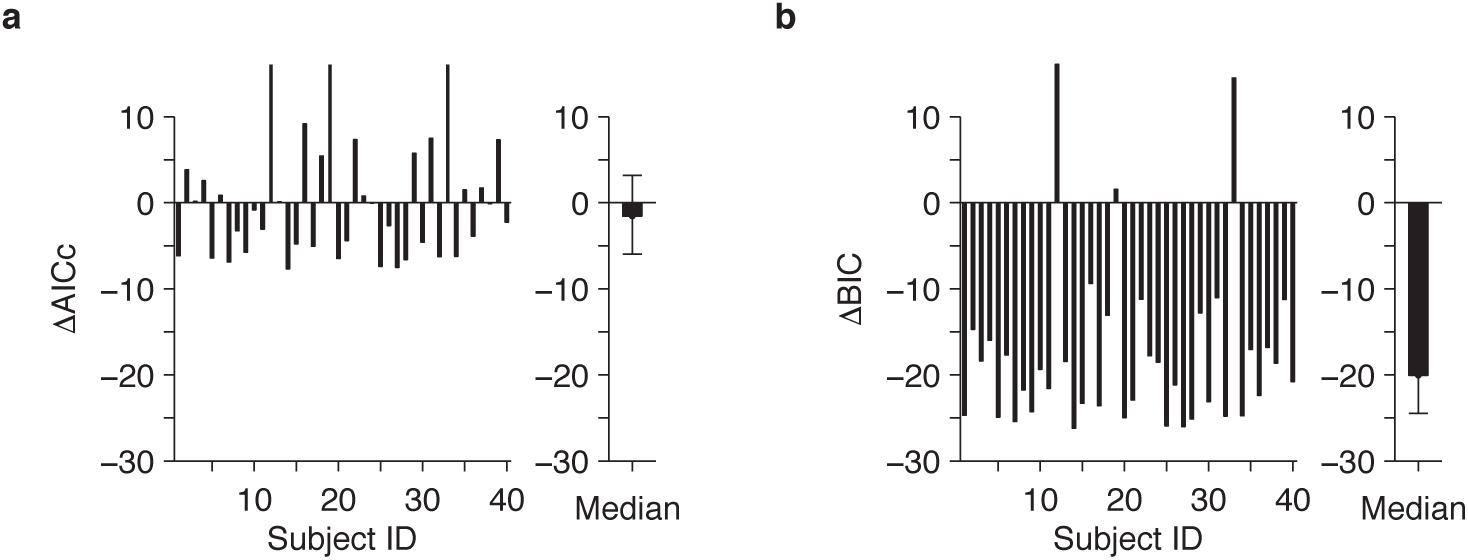
Model comparison justifies using the “shared” model. **(a)** AICc of the “shared” model minus AICc of the “full” model for **(Left)** individual subjects and **(Right)** Group - median and 95% bootstrapped confidence intervals. **(b)** Same for BIC.

We also performed three-way mixed-design ANOVA on the parameter estimates from the “full” model (Figure S8B). Just like in the “shared” model, we found a significant effect of group for log perceptual variability (*σ*) 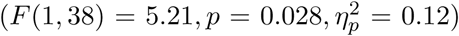, a significant effect of feature 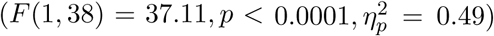, but no significant effect of load 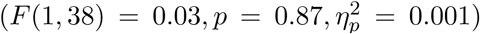. Neither of the two-way interactions nor the three-way interaction were significant (*p* > 0.06). After Sidak correction (*α* = 0.0065, 8 comparisons, since, as in the main model, we excluded across feature comparisons due to their different units) we found a between-group effect for Ori with *p* < 0.0001 and OriS (*p* < 0.0025), but no significant effects of group for neither Col (*p* = 0.0063) nor ColS (*p* = 0.47) (Figure S8).

For the log lapse *λ*, we found a significant effect of feature 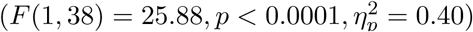 and a significant feature × group interaction 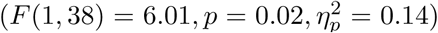; nothing else was significant (*p* > 0.09). After Sidak correction (*α* = 0.0043, all 12 comparisons make sense since *λ* is unitless), no between-group comparisons were significant (*p* > 0.02). Within Controls, the feature comparisons Ori vs Col and OriS vs ColS were significant (*p* < 0.001), but not the load ones. Within ADHD, neither the feature nor the load comparisons reached significance (*p* > 0.02).

For the PSE *µ*, like in the “shared” model, we found a significant effect of group 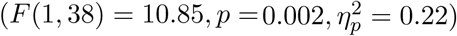 and also a significant group × load × feature interaction 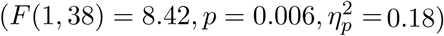, nothing else reaching significance (*p* > 0.09). After Sidak correction (*α* = 0.0065, as for *σ*), we found a significant difference between ADHD and Controls for ColS (*p* = 0.002), but not for Col (*p* = 0.18) and not for Ori or OriS (*p* > 0.12). Again, these results cannot provide robust support for ADHD participants having different S-cone dependent color processing.

**Figure S8:**
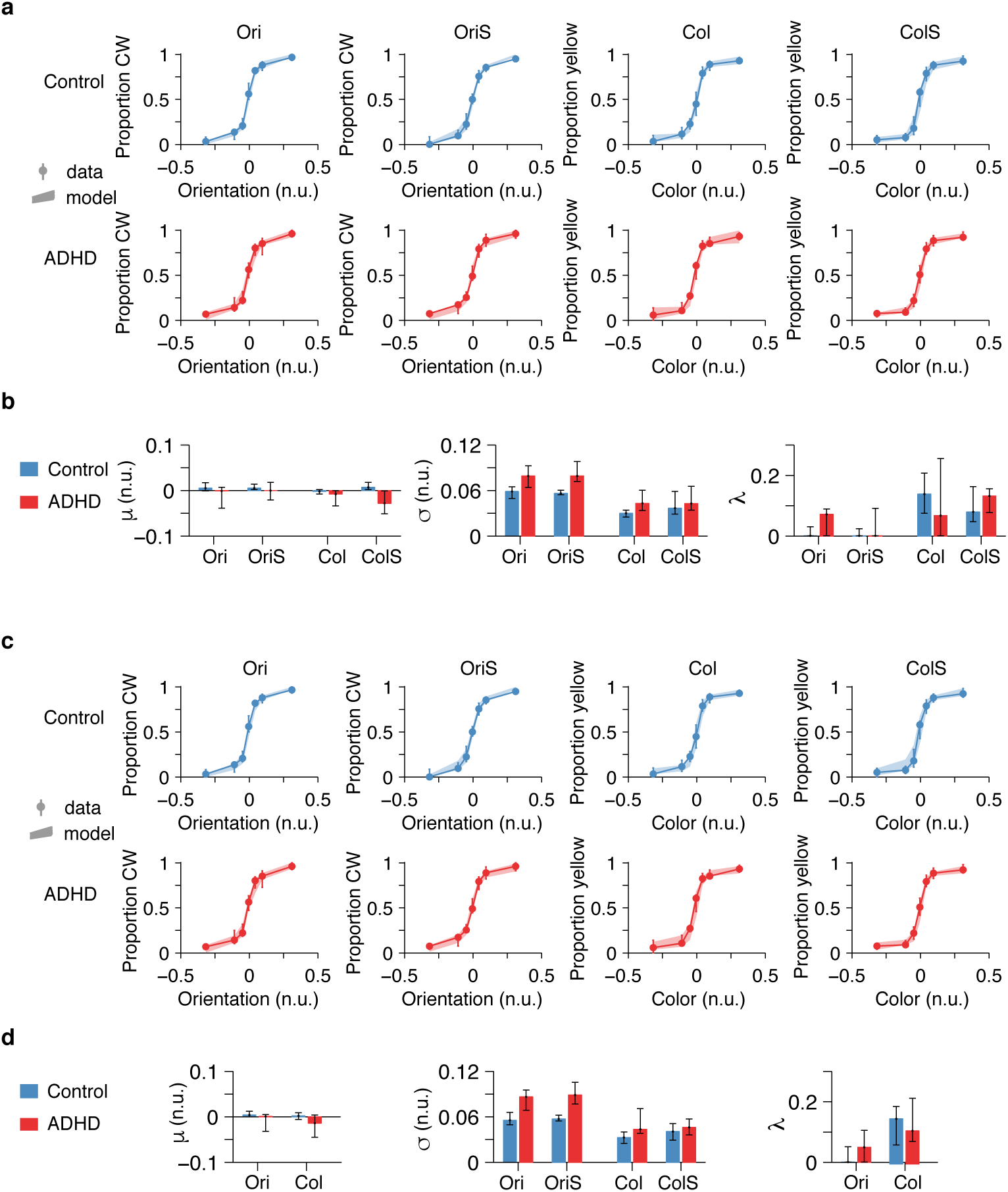
Psychometric curves for both models: data, model fits and parameter values. (a) “Full model” (12 parameters total): data and fitted psychometric curves. Solid circles with error bars show median and 95% bootstrapped confidence intervals, while shaded areas show the same for model predictions. The data was binned into 7 quantiles. Since the Bayesian adaptive method presented each participant in each condition with an unique set of stimuli, the midpoint stimulus values of the quantile bins differed for each. However, for ease of visualization, here we place the midpoints stimulus values for each bin as the midpoints obtained from binning into 7 quantiles the entire stimulus set concatenated across participants and conditions. **(b)** “Full model”: MLE parameter fits, **(c)** “Shared model” (8 parameters total): data and fitted psychometric curves. **(d)** “Shared model”: MLE parameter fits.

#### S4.4 Effect of learning

To assess learning across the experiment, we looked at the parameter estimates from the first half of the trials versus the second. Figure S9 shows that the perceptual variability parameters improved slightly on the second versus first half of trials, sign that there might be some learning. As reported in main, four-way mixed-design ANOVA on log *σ* confirmed a significant effect of time: 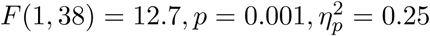. However, we note that these parameter estimates are not as reliable as the ones in Figure 4B, since they were obtained by fitting on only half the data.

**Figure S9:**
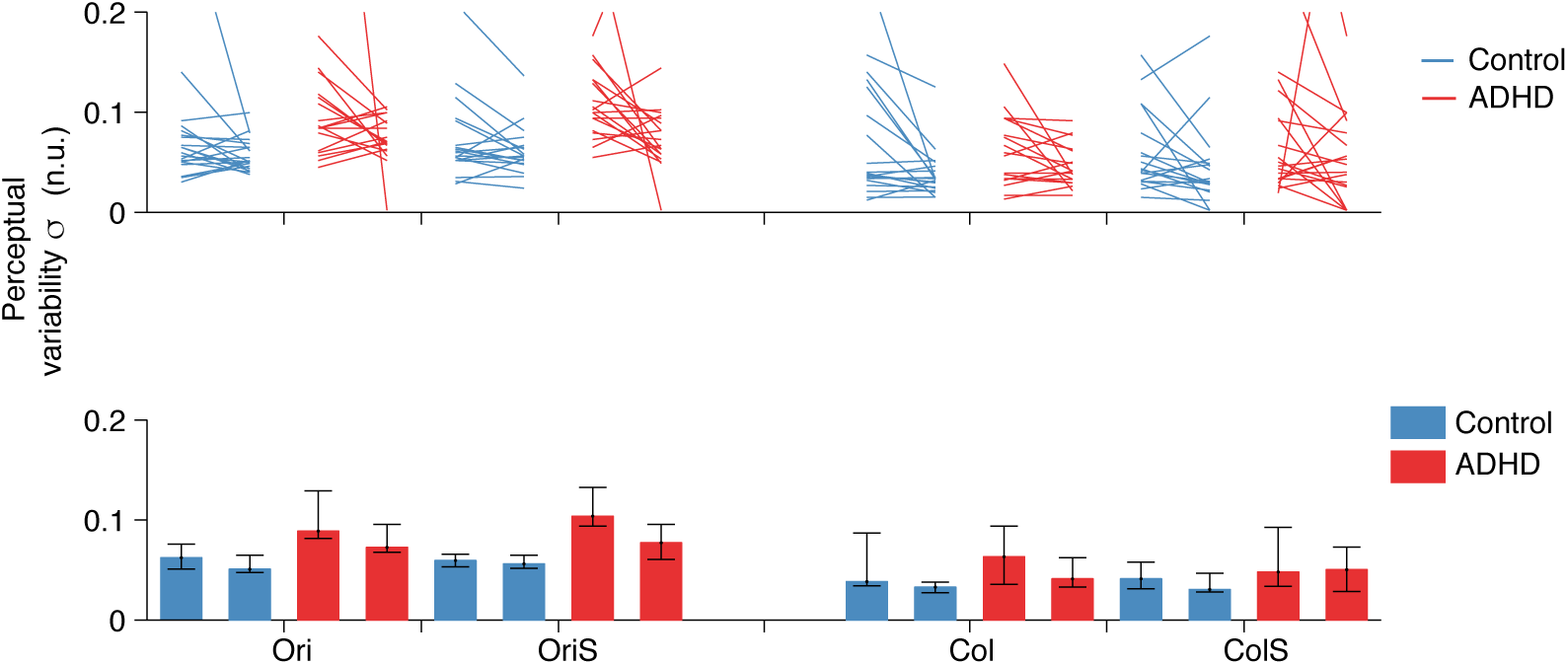
Perceptual variability parameter fits across time. **(Top)** Individual participants. Lines show how the perceptual variability differs from the first half of trials to the second half. **(Bottom)** Medians across participants and bootstrapped 95% confidence intervals.

### S5 Effect of eye tracking

A possible concern is that half of the participants in each group were eye-tracked, while half were not. If an eye-tracked participant broke fixation, they had to redo the trial. As a result, the eye-tracked participants started more trials (mean and SD for eye-tracked: 1047 ± 201 trials; non-eye-tracked always completed 800 trials). Thus, a concern could be that differences in task metrics could simply arise due to the experiment being longer and as a result more tiresome. We examined each of the average task metrics within each group, separately for the eye-tracked participants and the non-eye-tracked ones and found no significant differences (Wilcoxon rank-sum *p* > 0.13) (Figure S10).

**Figure S10:**
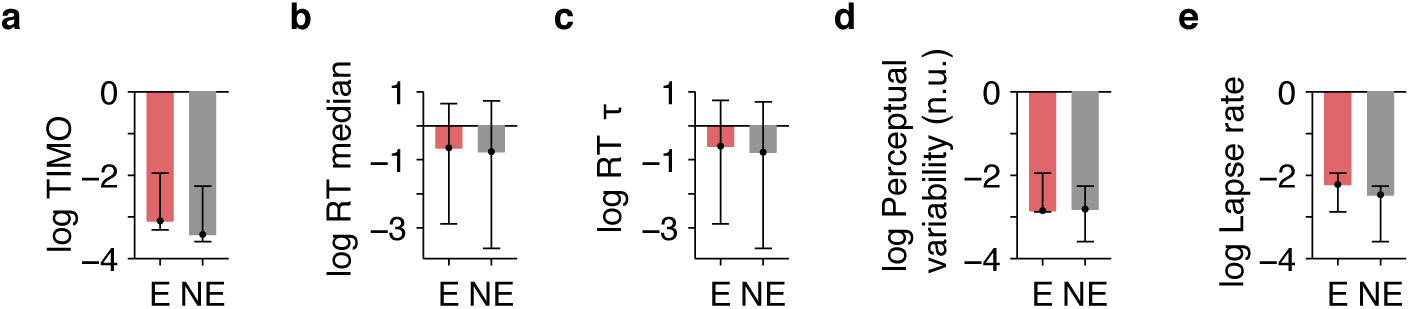
No significant difference between eye tracked (E) and non-eye tracked (NE) participants on behavioral task metrics: (a) TIMO, (b) RT median, (c) RT *τ*, (d) Perceptual variability, (e) Lapse rate. Bars represent medians and error bars bootstrapped 95% confidence intervals.

### S6 Breakdown of correlations

#### S6.1 By group

In Figure S11, we show the points that make up the correlations from Table 1, color coded by group. Of note, the two ADHD participants who had visibly lower orientation discrimination performance (Figure 4A), did not also have outstandingly reduced performance on other metrics; more detailed ophtamological examination could have provided more insight into the possible sources of their reduced orientation discrimination performance.

In Table S4 we show the pairwise correlations across task metrics separately within the Control group and within the ADHD group. Here, the only group specific correlations that survive the multiple-comparisons correction are RT with RT *τ* and GEC with ACDS.

In addition, we attempted to determine whether for a given pair of task metrics, their correlation within the ADHD group is different from their correlation within the Control group. To do this, we compared the difference between the actual correlations to a distribution of differences between correlations obtained by shuffling the ADHD and Control labels. We did not find significant differences between the ADHD and Control correlations for any pair of task metrics (*p* > 0.04).

**Figure S11:**
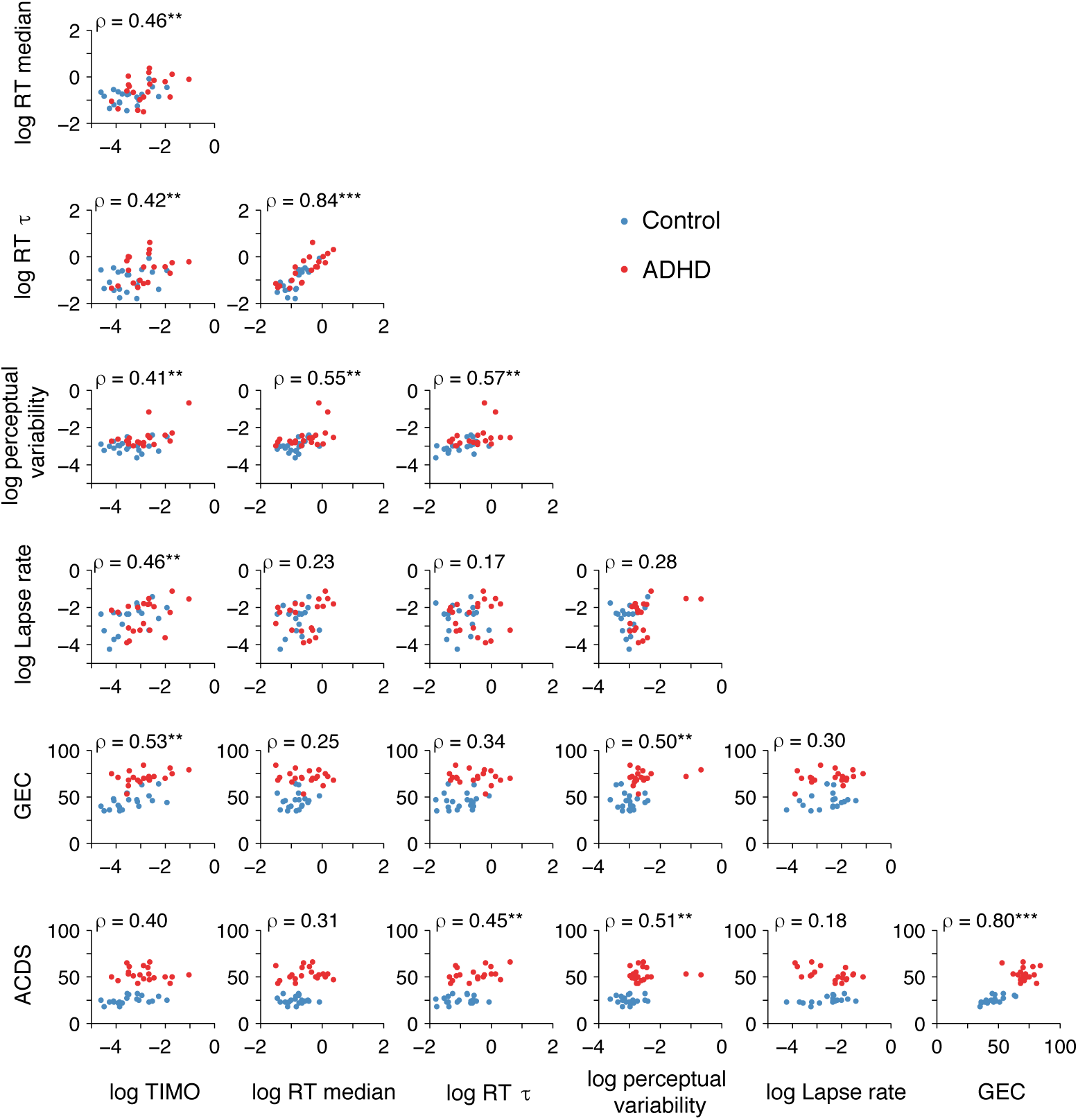
Dots: pairwise task metrics, color coded by group. We also show here the Spearman correlations collapsed across groups, as in Table 1. ** depicts 0.001 < *p* < 0.0089 (since *α*_Sidak_ = 0.0089 after multiple-comparisons correction) and *** depicts *p* < 0.001.

**Table S4:** No evident pattern of group specific correlations. Pairwise Spearman correlations across task metrics (both behavioral and clinical), as in Table 1, but divided by group. Boldfaced is significant after multiple-comparisons correction, *α* = 0.0083 for Control and *α* = 0.0082 for ADHD. (see Methods)

| (a) **Control** |  |  |  |  |  |  |
| --- | --- | --- | --- | --- | --- | --- |
| | TIMO | RT | RT $\tau$ | Perceptual variability ($\sigma$) | Lapse ($\lambda$) | GEC |
| TIMO |  |  |  |  |  |  |
| RT | $\rho = 0.25$<br>$p = 0.27$ | | | | | |
| RT $\tau$ | $\rho = 0.13$<br>$p = 0.59$ | **$\rho = 0.81$**<br>$p < 0.0001$ | | | | |
| Perceptual variability ($\sigma$) | $\rho = -0.02$<br>$p = 0.92$ | $\rho = 0.52$<br>$p = 0.02$ | $\rho = 0.50$<br>$p = 0.03$ | | | |
| Lapse rate ($\lambda$) | $\rho = 0.49$<br>$p = 0.03$ | $\rho = 0.22$<br>$p = 0.34$ | $\rho = 0.19$<br>$p = 0.42$ | $\rho = -0.05$<br>$p = 0.84$ | | |
| GEC | $\rho = 0.52$<br>$p = 0.02$ | $\rho = 0.16$<br>$p = 0.49$ | $\rho = 0.15$<br>$p = 0.52$ | $\rho = -0.12$<br>$p = 0.61$ | $\rho = 0.25$<br>$p = 0.29$ | |
| ACDS | $\rho = 0.49$,<br>$p = 0.03$ | $\rho = -0.09$<br>$p = 0.67$ | $\rho = 0.03$<br>$p = 0.89$ | $\rho = -0.09$<br>$p = 0.69$ | $\rho = 0.46$<br>$p = 0.04$ | **$\rho = 0.70$**<br>$p < 0.0001$ |
| (b) **ADHD** |  |  |  |  |  |  |
| | TIMO | RT | RT $\tau$ | Perceptual variability ($\sigma$) | Lapse ($\lambda$) | GEC |
| TIMO |  |  |  |  |  |  |
| RT | $\rho = 0.48$<br>$p = 0.03$ | | | | | |
| RT $\tau$ | $\rho = 0.36$<br>$p = 0.12$ | **$\rho = 0.82$**<br>$p < 0.0001$ | | | | |
| Perceptual variability ($\sigma$) | $\rho = 0.46$<br>$p = 0.04$ | $\rho = 0.52$<br>$p = 0.02$ | $\rho = 0.46$<br>$p = 0.04$ | | | |
| Lapse rate ($\lambda$) | $\rho = 0.40$<br>$p = 0.08$ | $\rho = 0.38$<br>$p = 0.10$ | $\rho = 0.12$<br>$p = 0.61$ | $\rho = 0.32$<br>$p = 0.16$ | | |
| GEC | $\rho = 0.37$<br>$p = 0.11$ | $\rho = -0.10$<br>$p = 0.68$ | $\rho = 0.19$<br>$p = 0.42$ | $r = 0.15$<br>$p = 0.54$ | $r = 0.18$<br>$p = 0.44$ | |
| ACDS | $\rho = -0.18$<br>$p = 0.45$ | $\rho = 0.14$<br>$p = 0.56$ | $\rho = 0.36$<br>$p = 0.11$ | $\rho = 0.03$<br>$p = 0.87$ | $\rho = -0.35$<br>$p = 0.12$ | $\rho = -0.09$<br>$p = 0.07$ |

#### S6.2 By symptom type

For this analysis, 2 ADHD participants were excluded due to missing AISRS records. A breakdown of the AISRS scores into inattentive and hyperactive shows that their correlations with task metrics recapitulate the correlations seen with ACDS. This is not unexpected, given the high correlation between ACDS and AISRS scores, as well as the fact that the AISRS inattentive and AISRS hyperactive scores were highly correlated (*r* = 0.89, *p* < 10^-13^).

#### S6.3 By condition

In Table 1, for each participant, we averaged each behavioral metric across all four conditions. In Figure S12, we present the correlations of perceptual variability with TIMO, RT and RT *τ* broken down by condition. The correlations that survived after multiple-comparisons correction are between perceptual variability (*σ*) and TIMO in the Ori condition, between *σ* and RT in the OriS and *σ* and RT *τ* in Ori and Col. Overall, we cannot conclude much from these patterns of results.

**Table S5:** No evident pattern of differential correlations by symptom type. AISRS inattentive and hyperactive correlations with behavioral task metrics are almost identical and largely recapitulate the ACDS correlations. Boldfaced represents *p* < 0.0089.

| | TIMO | RT | RT $\tau$ | Perceptual<br>variability ($\sigma$) | Lapse ($\lambda$) | GEC |
| --- | --- | --- | --- | --- | --- | --- |
| ACDS | $\rho = 0.40$<br>$p = 0.01$ | $\rho = 0.31$<br>$p = 0.05$ | $\rho = \mathbf{0.45}$<br>$p = 0.004$ | $\rho = \mathbf{0.51}$<br>$p = 0.0008$ | $\rho = 0.18$<br>$p = 0.26$ | $\rho = \mathbf{0.80}$<br>$p < 0.0001$ |
| AISRS<br>inattentive | $\rho = 0.41$<br>$p = 0.01$ | $\rho = \mathbf{0.43}$<br>$p = 0.006$ | $\rho = \mathbf{0.46}$<br>$p = 0.003$ | $\rho = \mathbf{0.51}$<br>$p = 0.001$ | $\rho = 0.22$<br>$p = 0.17$ | $\rho = \mathbf{0.65}$<br>$p < 0.0001$ |
| AISRS<br>hyperactive | $\rho = 0.41$<br>$p = 0.01$ | $\rho = \mathbf{0.48}$<br>$p = 0.002$ | $\rho = \mathbf{0.49}$<br>$p = 0.001$ | $\rho = \mathbf{0.63}$<br>$p < 0.0001$ | $\rho = 0.18$<br>$p = 0.27$ | $\rho = \mathbf{0.65}$<br>$p < 0.0001$ |

**Figure S12:**
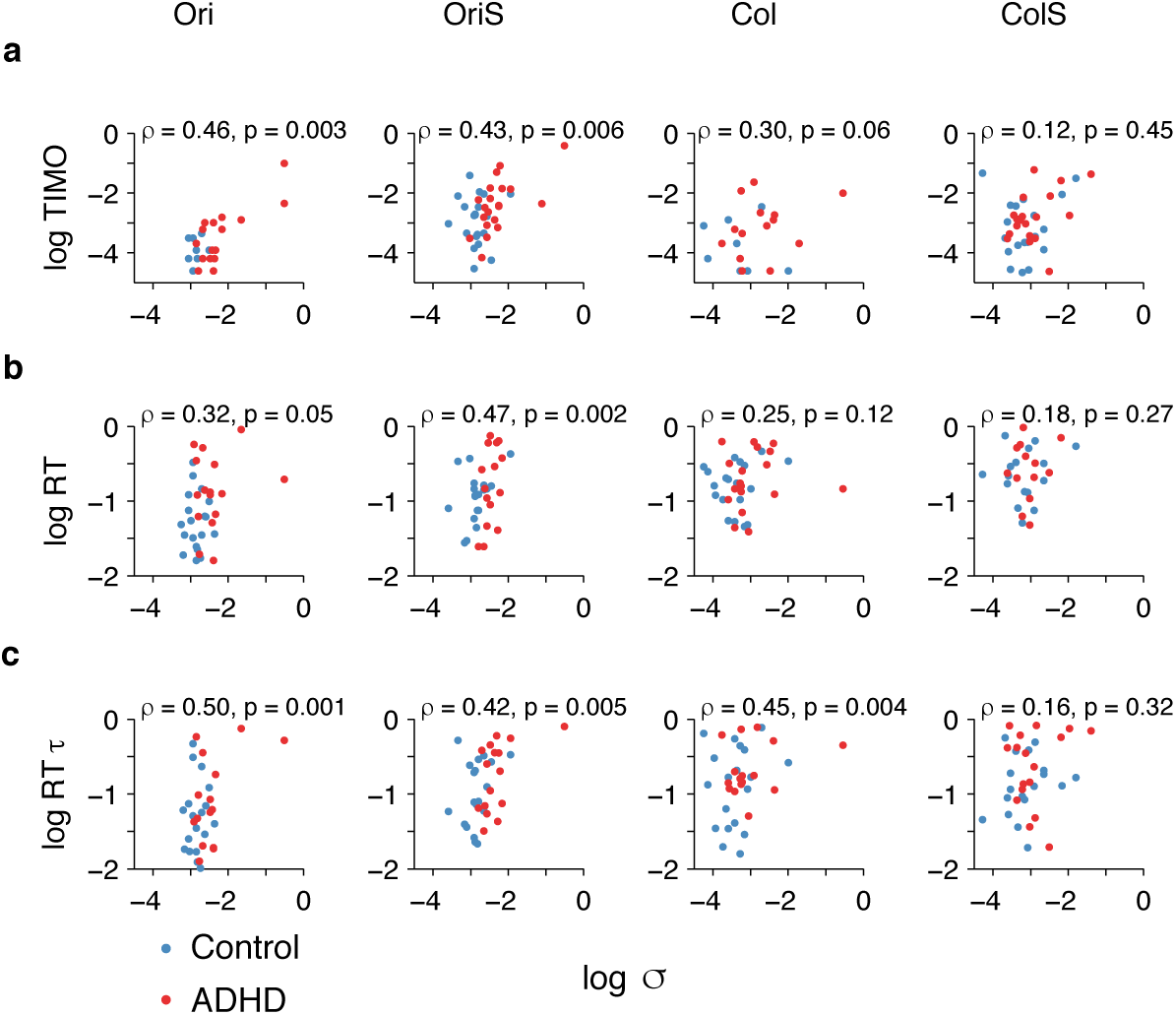
Spearman correlations of perceptual variability with other behavioral metrics broken down by conditions show no conclusive pattern. We show the correlations of log perceptual variability with (a) log TIMO, (b) log RT median and (c) log RT *τ*.

### S7 Prediction of clinical variables

#### S7.1 Logistic regression: prediction of diagnosis from behavioral metrics

**Table S6:** Logistic regression coefficients, mean ± sem.

| (a) **Diagnosis ∼ log perceptual variability** |  |  |  |
| --- | --- | --- | --- |
|  | coefficient | *t* value | *p* value |
| intercept | 13.3 ± 4.8 | 2.78 | 0.0055** |
| log perceptual variability ($\sigma$) | 4.7 ± 1.7 | 2.80 | 0.0051** |
| (b) **Diagnosis ∼ log TIMO** |  |  |  |
|  | coefficient | *t* value | *p* value |
| intercept | 1.9 ± 0.9 | 2.07 | 0.038* |
| log TIMO | 1.1 ± 0.5 | 2.25 | 0.024* |
| (c) **Diagnosis ∼ log perceptual variability + log TIMO** |  |  |  |
|  | coefficient | *t* value | *p* value |
| intercept | 13.3 ± 4.9 | 2.70 | 0.0069** |
| log perceptual variability ($\sigma$) | 4.3 ± 1.7 | 2.47 | 0.013* |
| log TIMO | 0.63 ± 0.59 | 1.07 | 0.28 |
| (d) **Diagnosis ∼ log perceptual variability + log TIMO + log RT median + log RT $\tau$ + log lapse rate** |  |  |  |
|  | coefficient | *t* value | *p* value |
| intercept | 14.0 ± 5.4 | 2.57 | 0.010* |
| log TIMO | 0.49 ± 0.64 | 0.76 | 0.44 |
| log RT median | -1.4 ± 1.7 | -0.81 | 0.42 |
| log RT $\tau$ | 1.4 ± 1.6 | 0.88 | 0.38 |
| log perceptual variability ($\sigma$) | 4.2 ± 1.9 | 2.23 | 0.025* |
| log lapse rate ($\lambda$) | 0.38 ± 0.37 | 1.02 | 0.30 |

### S7.2 Linear regression: prediction of clinical metrics GEC and ACDS from behavioral metrics

**Table S7:** Linear regression coefficients, depicted as mean ± sem, for GEC and ACDS with task metrics.

| (a) $\text{GEC} \sim \log \text{perceptual variability} + \log \text{TIMO} + \log \text{RT} + \log \text{RT } \tau + \log \text{lapse rate.}$ | | | | (b) $\text{ACDS} \sim \log \text{perceptual variability} + \log \text{TIMO} + \log \text{RT} + \log \text{RT } \tau + \log \text{lapse rate.}$ | | |
| --- | --- | --- | --- | --- | --- | --- |
| | coefficient | $t$ value | $p$ value | coefficient | $t$ value | $p$ value |
| log intercept | $103 \pm 12$ | 8.50 | $< 10^{-9***}$ | $73 \pm 13$ | 5.77 | $< 10^{-5***}$ |
| log TIMO | $8.0 \pm 3.2$ | 2.53 | 0.016* | $3.4 \pm 3.3$ | 1.04 | 0.31 |
| log RT median | $-10.3 \pm 7.6$ | -1.35 | 0.18 | $-11.3 \pm 8.0$ | -1.41 | 0.15 |
| log RT $\tau$ | $8.1 \pm 6.1$ | 1.32 | 0.19 | $14.4 \pm 6.4$ | 2.26 | 0.031* |
| log $\sigma$ | $6.1 \pm 4.9$ | 1.23 | 0.22 | $6.8 \pm 5.1$ | 1.33 | 0.19 |
| log $\lambda$ | $1.3 \pm 1.8$ | 0.75 | 0.46 | $0.5 \pm 1.9$ | 0.29 | 0.78 |

